# Labeled lines for fat and sugar reward combine to promote overeating

**DOI:** 10.1101/2022.08.09.503218

**Authors:** Molly McDougle, Alan de Araujo, Macarena Vergara, Mingxin Yang, Arashdeep Singh, Isadora Braga, Nikhil Urs, Brandon Warren, Guillaume de Lartigue

## Abstract

Food is a powerful natural reinforcer that guides feeding decisions. The vagus nerve conveys internal sensory information from the gut to the brain about nutritional value; however, the cellular and molecular basis of macronutrient-specific reward circuits are poorly understood. Here, we monitor *in vivo* calcium dynamics to provide direct evidence of independent vagal sensing pathways for detection of dietary fats and sugars. Using activity-dependent genetic capture of nutrient-specific vagal neurons activated in response gut infusions, we demonstrate the existence of separate hard-wired reward circuits for fat and sugar that are necessary and sufficient for nutrient-specific reinforcement. Even when controlling for calories, combined activation of fat and sugar circuits increases overeating compared to fat or sugar alone. This work provides new insight on the complex sensory circuitry that mediates motivated behavior and suggests that a subconscious internal drive to consume obesogenic diets (e.g., those high in both fat and sugar) may impede conscious dieting efforts.

## Introduction

The sharp rise in global obesity rates has been attributed to changes in the food environment that promotes overconsumption of palatable calorie-dense foods that are rich in fats and sugars ^1, 2^. Animal studies indicate that although palatability guides food choice, it is neither necessary ^3–5^ nor sufficient ^6^ to increase food intake. Instead, flavor preferences are rapidly learned as a conditioned response to post-ingestive nutritive cues ^7–10^. Thus, orosensory features of food are secondary to its nutritional value in underlying reinforcement ^11^. Importantly, the reinforcing value of intragastric infusions of fats or sugars enhance the overall intake of associated flavors ^12–14^. Thus, fats and sugars may cause overeating via a mechanism involving post-ingestive signaling, although the relative importance of individual macronutrients in diet-induced weight gain remains hotly debated ^1, 15–17^. We reasoned that defining the neural mechanisms for macronutrient-specific post-ingestive reward is of critical importance to understanding overeating.

The striatal regions of the basal ganglia are evolutionarily conserved brain substrates responsible for goal-directed behavior. Gustatory stimuli in the oral cavity recruit mesolimbic circuits that release dopamine in the ventral striatum, while nutritive stimuli that reach the intestine activate nigrostriatal dopamine release in the dorsal striatum (DS) ^18, 19^. Importantly, recruitment of the nigrostriatal circuit by post-ingestive stimuli occurs independently of palatability ^5, 18^, which presumably ensures prioritization of calories over flavor. Dopamine release from the substantia nigra pars compacta (SNc) onto the DS is critical for feeding. Genetic deletion of dopamine causes aphagia, yet normal eating behavior can be restored by reintroducing dopamine selectively in the DS ^20, 21^. Vagal sensory neurons have recently been identified as a key component of a multi-synaptic circuit linking the gut to nigrostriatal dopamine release ^22^. Optical stimulation of gut-innervating vagal sensory neurons is reinforcing, and animals actively seek stimuli that engage this circuit ^22^. Thus, animals will actively seek to consume nutrients sensed by vagal neurons that engage a nigrostriatal-dependent reward circuit. We aim to address whether fats and sugars recruit separate circuits.

Gastrointestinal infusion of fats ^23^ or sugars ^24^ activates the vagus nerve, and cause DS dopamine efflux ^25, 26^, yet previous attempts to address the role of the vagus nerve in appetition have resulted in mixed results ^25, 27–30^. Because vagal sensory neurons are molecularly heterogeneous ^31–33^ and play a role in diverse physiological functions ^34, 35^, we reasoned that previously used chemical or surgical lesioning approaches that interrupt global vagal activity lack the specificity required to test the role of distinct vagal sensory populations in nutrient reward. To overcome this issue we applied virally-delivered molecular tools to the nodose ganglia of FosTRAP mice to manipulate neuronal populations of the vagus nerve that are activated in response to gut detection of sugar or fat. Our results indicate that sugar and fat engage discrete neurons of the vaugs nerve and engage parallel but distinct reward circuits to increase feeding behavior.

### Fats and Sugars are Sensed by Distinct Vagal Populations

Fats ^23^ and sugars ^24^ both increase vagal firing in nerve recording experiments, however it remains unknown whether vagal neurons are generally activated in response to calories or if they sense specific macronutrients. To address this question we first adapted a technique for *in vivo* imaging of nodose ganglia neurons (Fig1A) ^33^ using mice expressing the Ca^2+^ indicator GCaMP6s driven by the pan-neuronal promoter Snap25 ^36, 37^. We used 2 photon microscopy coupled with an electrically tunable lens to enable recordings from nodose ganglion tissue in response to duodenal infusion of sugar (sucrose) or fat (corn oil) in the same live animal (Fig1A). A catheter was inserted into the duodenal bulb and an exit port created 2 cm from the pylorus to restrict delivery of macronutrient solutions to the proximal intestine (Fig1A). Analysis of individual neuronal responses indicated that spatially-defined vagal neuron subsets are activated in response to different nutrient stimuli (Fig1B-C). Nodose ganglia neurons exhibit either a strong response to fat, or a strong response to sugar, with few neurons responding to both (Fig1D-F). Notably, these intestinal nutrient-sensitive neurons accounted for a small fraction of the total population (<8%, Fig1D), supporting the concept of high cellular division of function within nodose ganglia ^32^. These data clearly indicate that fats and sugars recruit separate peripheral circuits.

**Figure 1.**
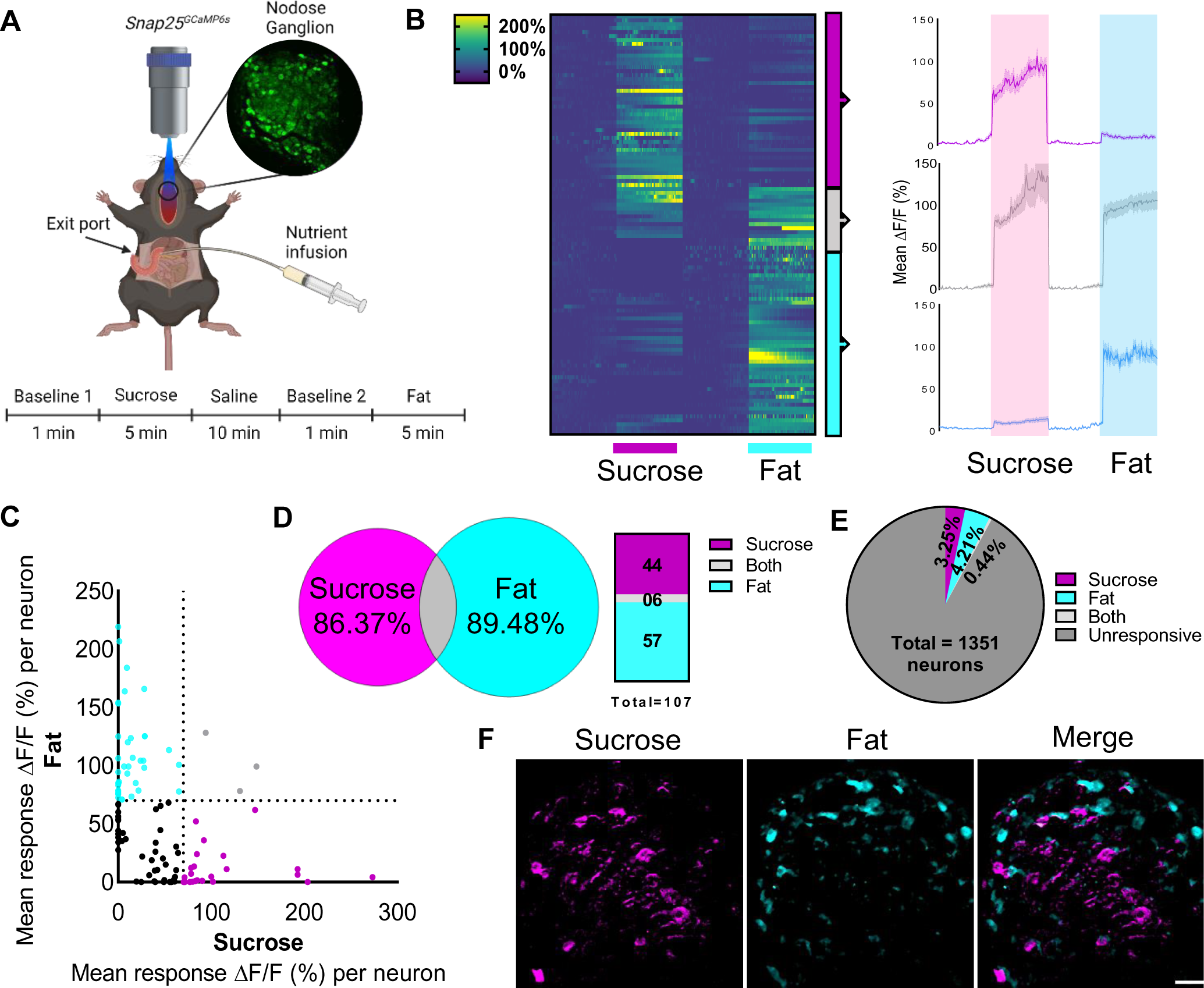
Sucrose and fat activate distinct vagal populations in the nodose ganglia. **A** In vivo imaging of the activity pattern of vagal sensory neurons in response to intraduodenal sucrose or fat infusions in *Snap25^GCaMP6s^*mice. **B** Heat maps depicting time-resolved responses (ΔF/F) of neurons identified as sucrose responders (magenta bar, 5 min) of fat responders (blue bar, 5 min); On the right, average ΔF/F of GCaMP6s signals in neurons that were responsive to sucrose (magenta), fat (blue) or both (gray). The shaded areas (magenta and blue) represent duration of stimuli. Dark lines represent means and lighter shaded areas represent SEM. **C** Average ΔF/F of GCaMP6s signal in neurons responsive to sucrose and/or fat; each dot represents a neuron that had at least one peak response of 70% or more above the baseline during the intraduodenal infusions; magenta dots = sucrose responsive; blue dots = fat responsive; gray dots = responsive to both stimuli; black dots = average response was below 70%. **D** Quantification of B. **E** Percentage of nodose ganglia neurons that were responsive to intraduodenal infusions of sucrose and fat. **F**. Image of the nodose ganglion of *Snap25G^CaMP6s^*mice showing calcium fluorescence responses to sucrose (magenta) and fat infusion (cyan). Scale bar, 100 µm. N=6.

### Macronutrient specific Reward

Next, we wanted to address the function of separate macronutrient-sensitive vagal sensory populations in reward. In order to genetically access NG neurons that were active in response to post-ingestive fat or sugar sensing, we used a transgenic mouse line that allows targeted recombination in active populations (TRAP2) ^38, 39^ (Fig 2A). These mice express an inducible cre recombinase, iCreER^T2^, under the control of an activity-dependent c-Fos promoter (*Fos^TRAP^* mice), enabling permanent genetic access to neuronal populations based on their activation to a defined, time-constrained stimulus when paired with injection of 4-hydroxytamoxifen (4-OHT) ^27, 40, 41^. We crossed the *Fos^TRAP^* mice with a Cre-dependent tdTomato reporter line, *Ai14* ^42^, and confirmed increased tdTomato^+^ nodose ganglia (NG) neurons in response to intragastric infusion of fat (intralipid, 7% w/v, Fat^TRAP^) or equicaloric sugar (sucrose, 15% w/v, Sugar^TRAP^) compared with iso-osmotic saline (Saline^TRAP^) (Fig2B-D) To validate the selectivity of the approach in NG neurons, we used Fos^TRAP^:Ai14 mice crossed with SNAP25^GCAMP6s^ mice and confirmed by live calcium imaging that responsive Sugar^TRAP^ NG neurons preferentially responded to intraduodenal infusions of sucrose (15% w/v) over isocaloric fat (SupFig1A,B). Furthermore, neurons of the nucleus tractus solitarius (NTS), that receive vagal inputs, had similar increased tdTomato labeling in response to intragastric infusion of fat or sugar compared to controls (Sup Fig1C-D). These findings are consistent with nutrient-induced Fos staining in the NTS ^43–45^, To validate the specificity of the Fos^TRAP^ approach to target macronutrient specific populations, we provide evidence that tdTomato labeling is dependent on 4-OHT (Sup Fig1D). Furthermore, we use cholecystokinin as a potent gastrointestinal stimulus known to require intact vagus nerve to mediate its feeding effects that can be repeatedly applied using IP injections. We confirmed that tdTomato-positive NTS neurons faithfully overlap with Fos-positive cells in response to CCK (Sup Fig2 A-C). By contrast, few tdTomato labeled CCK^TRAP^ NTS neurons were double labeled with Fos immunoreactivity after saline injection (Sup Fig2D-F). These results highlight the specificity and selectivity of the Fos^TRAP^ approach, and that we can efficiently and reliably gain genetic access to NG neurons that sense either fat or sugar.

**Figure 2.**
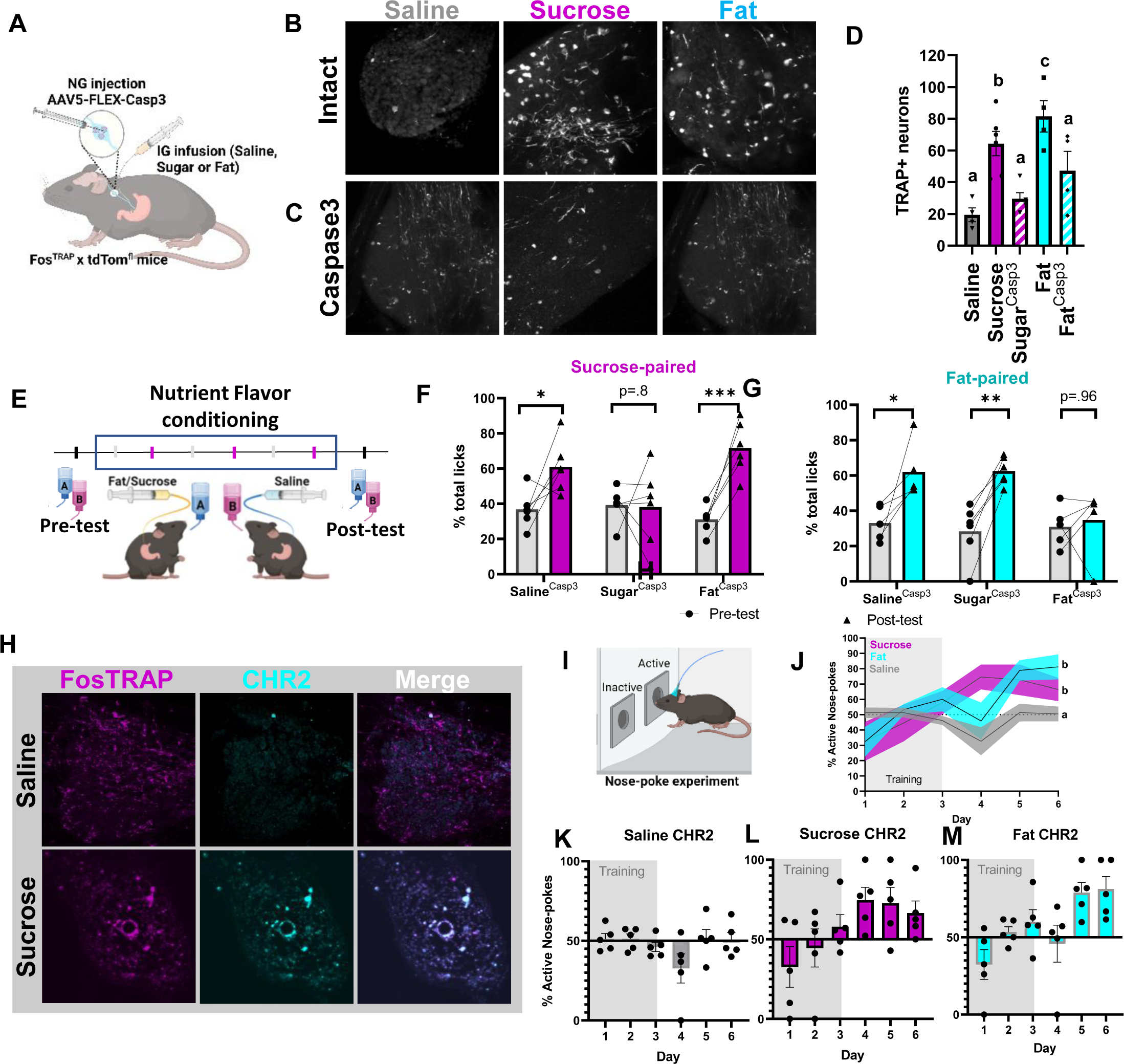
Sucrose and fat promote reward behavior via distinct vagal sensory populations. **A** Schematic of the FosTRAP approach to selectively ablate vagal sensory neurons that respond to intragastric infusion of fat or sugar. **B** FosTRAP targeting of nodose ganglia neurons increased tdTomato labeling in vagal sensory neurons in response to intragastric sucrose or fat infusion compared to saline. **C** Viral mediated ablation of vagal sensory neurons reduced tdTomato labeling to levels comparable to saline infusions. D Quantification of B and C n=4. One Way ANOVA with Tukey post hoc analysis.**E** Diagram demonstrating the flavor-nutrient conditioning paradigm. **F** Conditioning increases preference for the flavor paired with intragastic sucrose in Saline^TRAP^ and Fat^TRAP^ mice, but does not occur in Sugar^TRAP^ mice n=5-6. Two Way ANOVA with Bonferroni post hoc analysis **G** Conditioning increases preference for the flavor paired with intragastic fat in saline^TRAP^ and Sugar^TRAP^ mice, but does not occur in Fat^TRAP^ mice n=5-6. Two Way ANOVA with Bonferroni post hoc analysis **H** Representative images of ChR2 construct targeted to NG of Fos^TRAP^ mice. **I** Mice were trained in a nose-poke induced optogenetic self-stimulation paradigm. **J** After 3 days of training with food pellets in both nose holes, the mice in the Fat^TRAP^ and Sugar^TRAP^ groups favored the active nose hole. **K-L** Individual mouse data confirming that Saline^TRAP^ mice failed to learn to self-stimulate **(K)** while Sugar^TRAP^ **(L)** and Fat^TRAP^ **(M**) mice learned to nose poke for vagal stimulation n=5. Two Way ANOVA with Bonferroni post hoc analysis. Data are presented as mean ± s.e.m. \**P* < 0.05, \*\**P* < 0.01, \*\*\**P* < 0.001, NS, not significant.

Having validated this approach, we wanted to assess the necessity of separate macronutrient-sensitive vagal sensory populations for food reinforcement. As a measure of nutrient-driven behavioral reward we performed a flavor-nutrient conditioning task, in which animals are trained to prefer a novel flavor that has been experimentally-paired to intragastric infusion of nutrient (Fig2E). FosTRAP mice received bilateral NG injections of the viral construct AAV-flex-taCasp3-TEVp^46^ (Fig2C-D) which would enable selective ablation of vagal sensory neurons responsive to fat or sugar during the TRAP protocol. To confirm that viral injection of NG alone had no effect on post-ingestive reinforcement, we trained mice to associate flavors with either intragastric fat or sugar prior to TRAP. As expected, ^13, 14^ these mice were able to form a conditioned preference to each macronutrient (Sup Fig3A,B). Next, the TRAP protocol was performed to ablate the nutrient responsive populations. Mildly fasted Fos^TRAP^ mice were separated into 3 groups that received intragastric infusions (500 µl, 100 µl/min) of either saline, fat (microlipid, 7% w/v) or equicaloric sugar (sucrose, 15% w/v) followed by i.p. 4-OHT (30 µg/kg). This protocol deleted vagal sensory neurons based on their nutrient-responsive profile (Fig2D), while leaving other sensory and motor neurons intact. Control Saline^TRAP^ mice still formed robust preferences to novel, non-nutritive flavors paired with either intragastric infusion of fat or sugar (Fig2F-G). Fat^TRAP^ mice with deletion of fat sensing NG neurons formed normal preference for the flavor paired with sugar, but failed to reinforce the flavor paired with fat (Fig2F-G). Conversely, Sugar^TRAP^ mice with deletion of sucrose sensing NG neurons had selectively abolished sugar reinforcement (Fig2F-G). These striking findings demonstrate that separate, mutually exclusive populations of vagal neurons independently drive fat or sugar reward.

To assess whether stimulation of fat or sugar responsive vagal neurons are each independently sufficient for reward, mice underwent a previously-validated self-stimulation behavioral task ^22^ in which optogenetic stimulation of vagal sensory terminals is paired to nose poke (Fig2I). Fos^TRAP^ mice received bilateral NG injection of the *Cre*-inducible viral construct AAV9-EF1a-DIO-hChR2(H134R)-EYFP (ChR2) ^47^ to selectively express the light-sensitive depolarizing channel *Channelrhodopsin-2* (ChR2) in vagal afferent neurons (Fig2H). An optic fiber was placed above vagal terminals in the medial NTS ^22, 48^, and ChR2 was trapped in NG neurons in response to intragastric fat, sugar or saline. Optogenetic activation of vagal terminals increased Fos expression throughout the NTS compared to control mice that lack ChR2 (Sup Fig4A,B). Saline^TRAP^ mice (Fig2K) did not learn to prefer the nose hole that triggers optogenetic stimulation of vagal terminals. In contrast, both Sugar^TRAP^ (Fig2L) and Fat^TRAP^ (Fig2M) mice learned to self-stimulate vagal terminals, a hallmark behavior of reward ^49, 50^. Together, these findings reveal that separate post-ingestive sensing of fat and sugar are both necessary and sufficient for the development of macronutrient-specific reward.

### Fat and sugar responsive NG neurons innervate different organs

Based on the evidence that fats and sugars are signaled from different vagal populations, we next wanted to determine the site of fat and sugar reward. We used genetic guided anatomical tracing of fat and sugar responsive vagal afferents by injecting the cre-dependent virus AAVPhp.s-Flex-tdTomato bilaterally into NG of Fos^TRAP^ mice (Fig3A). Striking differences in peripheral innervation patterns were observed between fat and sugar sensing vagal populations. In the duodenum, we observed extensive innervation of mucosal endings in intestinal villi from Fat^TRAP^ mice and Sugar^TRAP^ mice but not Saline^TRAP^ mice (Figure 3B). Furthermore, we observed abundant vagal-derived *tdTomato^+^* fibers running along the portal blood vessels from Sugar^TRAP^ mice (Fig3C), but not in Fat^TRAP^ or Saline^TRAP^ mice. These data suggest that the duodenum is a key sensing site for both fat and sugar sensing, but sugar sensing also occurs at the level of the hepatic portal vein (HPV).

**Figure 3.**
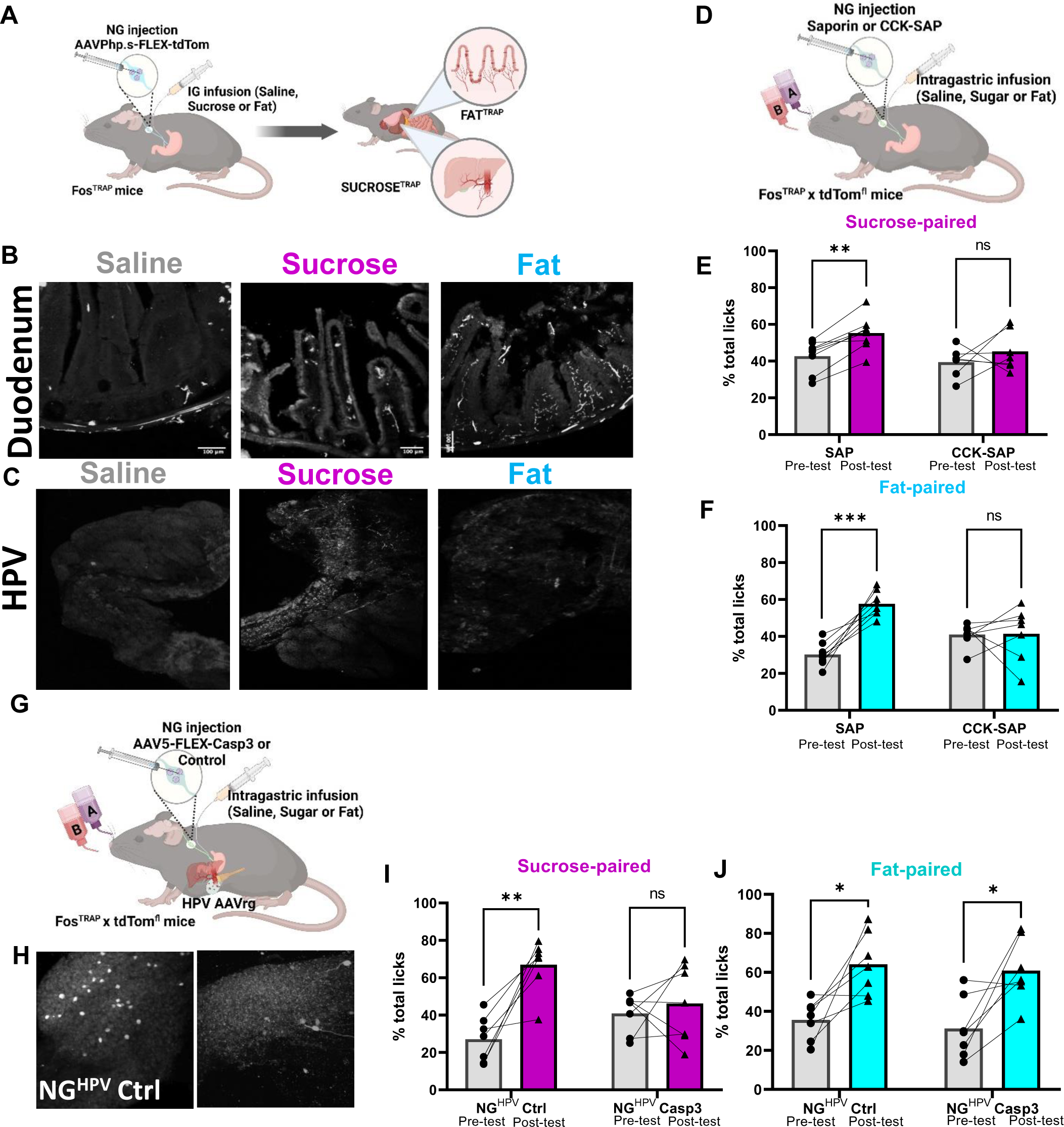
Anatomical dissociation of peripheral projections of sucrose and fat responsive NG neurons A. Schematic of viral guided mapping of vagal sensory neuron projections by injecting AAV-DIO-tdTomato virus bilaterally into NG of FosTRAP mice in response to intragastric saline, sucrose (15%), or fat (7%). **B** Terminals visualized by immunofluorescence of the duodenum are most abundant in Fat^TRAP^ mice. N=4 **C** The hepatic portal vein exclusively receives innervation from Sugar^TRAP^ mice. N-4 Two Way ANOVA with Bonferroni post hoc analysis **D** Schematic of approach for vagal deafferentation of the gut using CCK-SAP. **E** CCK SAP treatment abolishes flavor conditioning in response to intragastric infusion sucrose. N=7-8. Two Way ANOVA with Bonferroni post hoc analysis. **F** CCK SAP treatment abolishes flavor conditioning in response to intragastric infusion fat.n=7-8. Two Way ANOVA with Bonferroni post hoc analysis. **G**.Combinatorial viral approach for vagal deafferentation of the HPV. **H** Representative images of labeling in a subsets of NG neurons innervating the HPV can be selectively ablated. **I-J** Loss of NG^HPV^ neurons abolishes sucrose (i) but not fat (j) reinforcement. N=7. Two Way ANOVA with Bonferroni post hoc analysis. Data are presented as mean ± s.e.m. \**P* < 0.05, \*\**P* < 0.01, \*\*\**P* < 0.001, NS, not significant.

To assess the importance of the gastrointestinal vagal fibers for fat signaling we injected the neurotoxin saporin conjugated to cholecystokinin (CCK-SAP) bilaterally into the NG of WT mice (Fig3D, SupFig5A) a previously validated method for selective vagal deafferentation of the upper GI tract ^51^. We find that mice with confirmed CCK-SAP deafferentation (Sup Fig5B) significantly increased oral consumption of 7.5% fat solution (Sup Fig5C), but this produced no change on the intake of equicaloric sucrose solution (Sup Fig5D). These data are consistent with evidence that vagal deafferentation of the gut abolishes fat, but not sugar, satiety in rats ^52^; instead, sugars inhibit hunger via a spinal gut-brain circuit ^53^. As predicted from the viral tracing experiments, vagal deafferentation of the upper gut using CCK-SAP treatment blocked conditioned preference for both fat and sugar (Fig3E, F). These data support prior work showing that CCKR is expressed in many different mechanosensory and chemosensory vagal sensory subpopulations ^32^ that innervate the length of the GI tract including the HPV ^31^. Thus, CCKR is a viable molecular marker to dissociate vagal sensory populations that convey fat and sugar satiety, but not post-ingestive reward.

Our results revealed vagal innervation of the HPV by SugarTRAP neurons. These data are consistent with previous work showing that the HPV receives nutrient-rich blood from multiple gastrointestinal organs ^54^, and vagal sensory innervation of the HPV wall is implicated in sensing of dietary glucose ^55^ and food preference ^56, 57^. To test the role of HPV innervating vagal sensory neurons in post-ingestive reward, we adapted a technique for targeting NG neurons based on their pattern of innervation ^22^ (Fig3G). We first confirmed that a subset of NG neurons was labeled in response to brushing Cre-expressing retrograde adeno-associated virus (AAVrg-cre) on the HPV wall of Ai14 mice (Fig3H). Next, we selectively ablated HPV-innervating NG neurons, by bilaterally injecting cre-dependent caspase virus in the NG and applying AAVrg-Cre to the HPV wall of Ai14 mice (Fig3G, H). In flavor-nutrient conditioning experiments, we found that deletion of HPV-innervating NG neurons abolished sugar (Fig3I), but not fat (Fig3J), reinforcement. Thus, we provide anatomical and behavioral data supporting a necessary role for HPV innervating vagal sensory neurons in post-ingestive sugar reward. These data support the idea that fat and sugar reward are mediated by separate vagal sensory populations that can be distinguished based on their peripheral innervation patterns.

### “Labeled lines” for fat and sugar reward

The vagus nerve acts as key neural relay that connects the gut to dopamine producing SNc neurons ^22^. Optogenetic activation of gut-innervating vagal sensory neurons results in dopamine efflux in the DS and produces reward behaviors ^22^. Thus, gut-innervating vagal sensory neurons are capable of mediating reward to post-ingestive nutritive signals. Consistent with this notion, <u>microdialysis</u> sampling from the DS (Fig4A) in mildly food-restricted awake mice results in rapid and sustained dopamine efflux in response to intragastric infusion of fat or sugar independently (Fig 4B).

**Figure 4.**
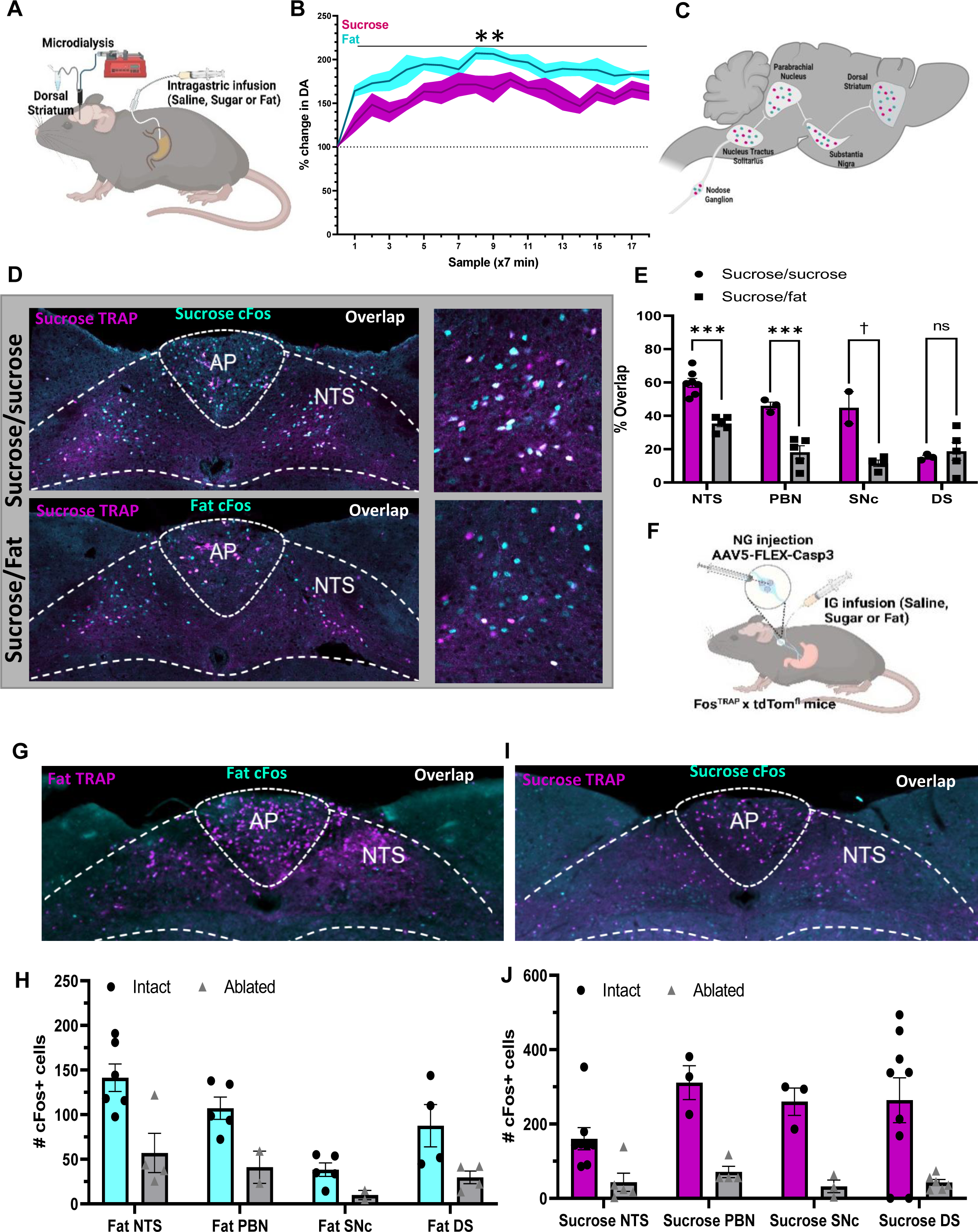
Fat and sugar recruit parallel but separate reward circuits. **A** Schematic illustrating microdialysis of the dorsal striatum performed to measure dopamine in response to 5 minutes of intragastric sucrose (15%, 100ul/min) or fat (7%) compared to saline baseline. **B** Rapid and sustained dopamine levels were found in response to both intragastric infusion of fat or sugar. N=5. Two Way ANOVA with Bonferroni post hoc analysis**. C** Schematic comparing neuronal activity in response to IG sucrose and IG fat along the gut-reward circuit. **D** Representative images of the NTS in response to sucrose (TRAP, magenta), and either high colocalization after infusion two weeks later of sucrose (Fos, cyan; top) or low colocalization after infusion of fat (Fos, cyan; bottom) in the same animal. Higher magnification images on the right. **E** Quantification showing higher overlap between repeated infusions sucrose compared to separate macronutrients in the NTS, PBN, and SNc. N=3-7. Unpaired 2 way Student’s t-test. **F** Schematic representing mice that received caspase mediated ablation of NG neurons in response to intragastric infusion of nutrients to test the necessity of these neurons in recruiting fat or sugar specific gut-reward circuitry. **G** Representative images of brain activity in response to fat was compared in the same mouse before (TRAP) and after (cFos) nutrient induced ablation of NG neurons. **H** Ablation of fat sensing NG neurons blunts the response to subsequent response to fat at each level of the gut-reward circuit. **I** Representative image of brain activity in response to sucrose was compared in the same mouse before (TRAP) and after (cFos) ablation of sucrose sensing NG neurons. N=3-6. Unpaired 2 way Student’s t-test **J** Ablation of sugar sensing NG neurons blunts the response to subsequent response to sugar along the entire length of the gut-reward circuit. N=3-6. Unpaired 2 way Student’s t-test Data are presented as mean ± s.e.m. \**P* < 0.05, \*\**P* < 0.01, \*\*\**P* < 0.001, NS, not significant.

Having demonstrated that fats and sugars are both capable of activating a nigrostriatal dopamine circuit, we inquired whether post-ingestive signals diverge at the cellular level along the well-defined gut reward circuit ^22^ that comprises the NTS➔ dorsolateral Parabrachial nucleus (dlPBN) ➔ SNc ➔ DS (Fig4C). We used Fos^TRAP^ mice to address the challenge of comparing neuronal activity in response to multiple stimuli at different timepoints over multiple brain regions in the same mouse. We validated the ability to compare Fos^TRAP^ neurons labeled with tdTomato and IF-labeled cFos-expressing neurons following exposure to the same or different stimuli separated by 14 days in the same animal (Fig4D, and Sup Fig6A-C). These findings support the concept that unique neuronal ensembles encode macronutrient-specific post-ingestive reward. When analyzing the overlap between sucrose TRAP and sucrose Fos labeling, we found high overlap in the NTS, PBN, and SNc (Fig4E). However, when comparing sucrose TRAP and fat Fos labeling, there was only limited overlap between the neurons labeled during the two separate nutritive stimuli (Fig4E). Specifically, the overlap was reduced by half between intragastric infusions of equicaloric sucrose and fat in the NTS and dlPBN, and reduced by half again at the level of the SNc, suggesting greater differentiation of nutrient signaling at distal nodes of the gut reward circuit (Fig4C,D).

There was low overlap between sugar TRAP and sugar Fos in the DS, suggesting that the DS integrates dopamine release independently the type of nutritive signal that causes the release. Importantly, we observed high overlap of FosTRAP and Fos IF neurons in response to separate intragastric stimuli in the PVT and VTA (Sup Fig6D-G), suggesting that these brain regions are more attuned to the availability of food or physiological state than to the specific macronutrient composition of a meal. Together, these data indicated that fats and sugars recruit parallel and largely separate neuronal populations at each node of the gut-reward circuit, supporting the existence of labeled lines for fat and sugar reward.

Using the same approach, we addressed the necessity of nutrient-sensitive vagal sensory neurons in recruiting the nigrostriatal system. We compared within the same animal, the neuronal activity along the gut-nigrostriatal circuit in response a nutrient before and after deletion of vagal sensory neurons (Fig4F). Cre-dependent caspase virus was bilaterally injected into the NG of FosTRAP mice, and neurons were trapped in response to intragastric infusion of sugar or fat with 4-OHT. This approach resulted in TRAP labeling with tdTomato in responsive neurons (SupFig7A and B), while the caspase virus resulted in targeted ablation of vagal sensory neurons that respond to the stimuli (SupFig7A and B). Robust tdTomato labeling was observed throughout the brain in response to both fat and sugar infusions (SupFig7A and B), indicating that the caspase virus had little impact on central processing of post-ingestive signals. However, Fos IF was significantly blunted in the NTS (Fig4G-J), PBN (Fig 4H, J and SupFig7A-B), SNpc (Fig 4H, J and SupFig7A-B), and DS (Fig 4H, J and SupFig7A-B) in response to individual macronutrients. These data suggest that chemosensory vagal neurons are a necessary relay for post-ingestive reward for both fat and sugar.

### Combination of fat and sugar promote overeating for pleasure

The implication of the above data is that combined activation of fat and sugar reward circuits may be more rewarding and appetitive than fats or sugars alone. To test this hypothesis, we quantified 30 min *ad libitum* intake in WT mice using a contact lickometer as mice voluntarily licked equicaloric solutions of fat, sucrose, or a mixture of the two macronutrients (Fig5A). Mice readily consumed similar amounts, and strikingly, mice nearly doubled the number of licks for the solution containing a combination of fat and sugar compared to solutions containing fat or sugar alone (Fig5B). We next assessed if mice show the same pattern of intake if the nutritive aspects of the solution are isolated (e.g., if we remove flavor). Food-restricted WT mice with surgically-implanted gastric catheters licked an non-nutritive flavored solution paired with intragastric infusions of equicaloric solutions of sugar, fat, or the combination (Fig5C).

**Figure 5.**
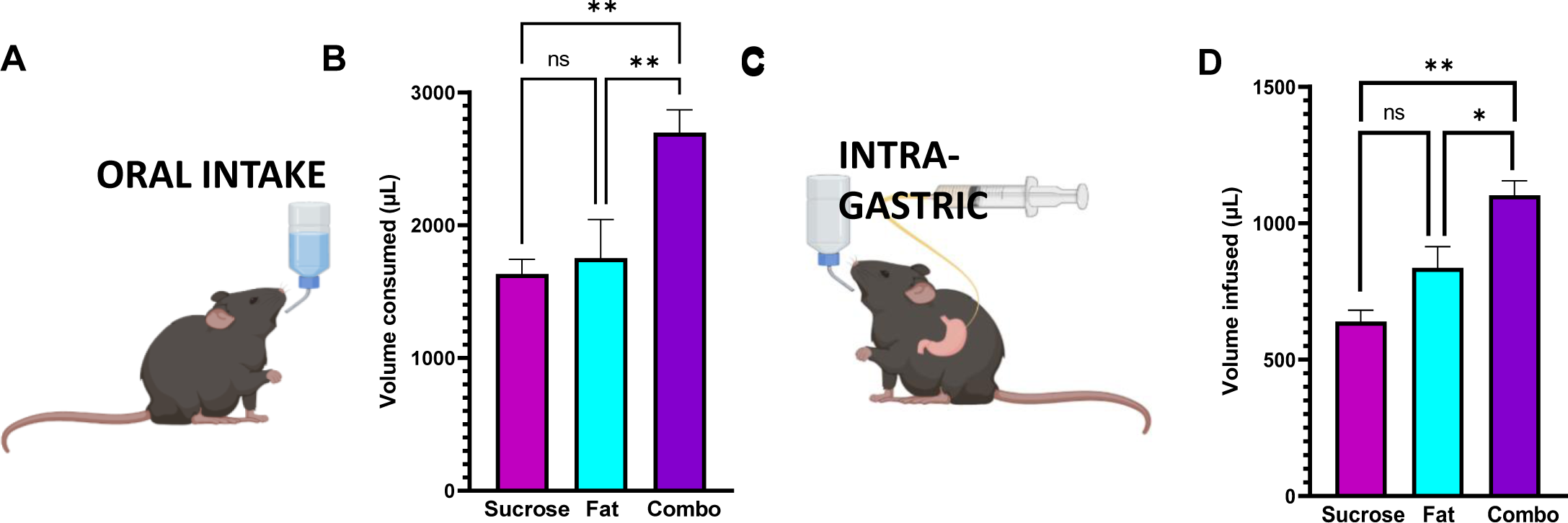
Combining fat and sugar additively increases intake and reward signaling. A. Contact lickometer were used to measure mice voluntary intake of equicaloric solutions of fat, sugar or a mixture containing fat and sugar. **B** Mice consumed larger volume of equicaloric combined solution than either fat or sugar alone in a one bottle licking test. n=15. One Way ANOVA with Tukey post hoc analysis **C** Mice implanted with intragastric catheters licked non-nutritive flavored solutions for intragastric infusions of equicaloric solutions of sugar, fat or a solution containing both fat and sugar. **D** Mice licked more to receive intragastric infusion of solution containing a combination of fat and sugar compared to equicaloric solutions of fat or sugar alone. n=15. One Way ANOVA with Tukey post hoc analysis. Data are presented as mean ± s.e.m. \**P* < 0.05, \*\**P* < 0.01, \*\*\**P* < 0.001, NS, not significant.

After training, the mice exhibited the same elevated licking pattern for the mixed solution in this experimental paradigm(Fig5D), suggesting that the increased preference for the combination of fat and sugar is independent of taste and is the result of post-ingestive signaling.

## Discussion

Our findings reveal the existence of labeled lines for fat and sugar reward. Using in vivo calcium imaging of nodose ganglia we first identify two vagal sensory populations that respond almost exclusively to post-ingestive fat or sugar, and then employ a Fos^TRAP^ approach to delineate the anatomy and function of these neurons. We reveal that these two distinct vagal populations 1) innervate separate peripheral organs, 2) are necessary for nutrient specific reinforcement, and 3) initiate parallel gut-brain reward circuits that can be dissociated at the cellular level. The activation of separate fat or sugar circuits can increase motivated feeding behavior, but strikingly, the combination of fats and sugars potentiate overeating independently of calories. Thus, we describe how parallel activation of food reward circuits transmits the internal drive to consume obesogenic foods.

### Identifying the function of chemosensory NG neurons

Our data identify a novel role of chemosensory NG populations in signaling nutrient reward. The nodose ganglia is composed of heterogenous populations of vagal neurons responsive to various sensory modalities. Two populations of tension-sensitive mechanosensory NG neurons have been identified that form either intramuscular arrays in the muscle layers or intraganglionic laminar endings in the myenteric plexis ^34, 58^. A separate chemosensory NG population was identified in the 1950’s ^59^ based on electrophysiological responses to chemical, but not mechanical, stimuli ^59–61^. Tracing experiments supported that these chemosensory NG neurons form distinct mucosal endings in intestinal villi ^31^ and suggested that they were ideally situated to respond to absorbed nutrients and hormone products from the gut epithelium. Transcriptomic sequencing of NG later confirmed that chemosensory populations are genetically distinct from mechanosensory neurons ^31^. Although optogenetic stimulation of genetically-defined mechanoreceptors potently inhibit acute food intake, chemosensory neurons did not; thus the function of chemosensory neurons remains elusive ^31^. Here we used Fos^TRAP^ to functionally target a population of NG neurons that are nutrient-sensitive and demonstrate that these neurons form mucosal terminals in intestinal villi, hallmarks of chemosensory neurons. Using viral mediated ablation and optogenetic stimulation approaches we demonstrate that these chemosensory neurons engage hardwired gut-brain reward circuits and cause nutrient-specific reinforcement and appetition. Several prior studies using less precise tools including resection of subdiaphragmatic vagus nerve have resulted in variable outcomes ^29, 30, 62^. These types of experiments are difficult to interpret because of the confound caused by the deletion of vagal motor signaling that controls gut function. This issue is partially addressed by chemically-induced vagal deafferentation using capsaicin ^63^, yet capsaicin experiments also show no impairment of fat or sugar reinforcement ^28^. Notably, capsaicin causes cell death by acting at TRPV1 channels which are expressed at very low levels in many chemosensory NG neurons ^32^ suggesting capsaicin treatment may spare key vagal sensory neurons required for fat and sugar reinforcement. Importantly, more selective approaches, that do not impair vagal control of gut function, support a role for the vagus nerve in relaying reward information to the brain ^22, 27, 64^.

Using a Fos^TRAP^ strategy that enables the most highly-targeted nutrient-specific vagal deafferentation to date, we demonstrate that separate vagal circuits exist to mediate fat and sugar reinforcement. We provide multiple lines of evidence that FosTRAP is an effective strategy that provides permanent genetic access to vagal sensory neuron populations based on their response to defined sensory stimuli, unbiased by genetically-defined cell type. First, using calcium imaging we confirm that Sugar^TRAP^ NG neurons are activated in response to intraduodenal infusion of sugar, but not fat. Second, we demonstrate that caspase-mediated ablation of nutrient responsive NG neurons prevents nutrient-specific reinforcement and third we show that ablation of nutrient-specific vagal circuit impairs downstream neural activity in the brain. Finally, we demonstrate that tracing from Fat^TRAP^ and Sugar^TRAP^ result in different gastrointestinal innervation patterns. These data support prior work predicting genetic heterogeneity based on sensory modality ^32^, but suggests a previously unsuspected amount of cellular diversification according to sensitivity to individual luminal nutritive stimuli. Thus, we provide direct evidence that chemosensory neurons play a key role in food choice that promotes survival by ensuring consumption of nutritive foods.

According to our data, vagal sensory neurons that express CCK receptor are necessary for both fat and sugar reinforcement, but these separate populations can be further subcategorized based on their innervation of the HPV. The lack of specificity of CCK receptor as a marker for fat or sugar is not surprising given their abundance in the NG. Prior sequencing data of vagal sensory neurons identified multiple clusters of *Cckar* neurons, including a chemosensory cluster t6 ^31^ or G ^32^ characterized by co-expression of *Vip*, *Htr3a* and *Ust2b*, and a separate polymodal cluster t3/t1 ^31^ or I ^32^ that co-express *Oxtr*, *Car8*, and *Ctnx2*. Response to luminal infusion of glucose was detected by both of these clusters ^32^ and a fraction of neurons in each of these clusters innervate the HPV and the duodenum ^31^. Another large population of chemosensory neurons characterized by *Gpr65* expression were identified in cluster t9-12 ^31^ and F ^32^. The role of NG*^Gpr^*^65^ neurons as fat and/or sugar sensors remains unclear because 1) they do not appear to co-express *Cckar* and 2) prior work suggests that they exhibit broad response to intestinal stimuli ^33^ consistent with a role as polymodal osmosensors ^65^. Thus, fat and sugar NG neurons may be comprised of genetically-heterogeneous populations and it remains unclear whether they can be delineated on the basis of a single genetic marker. An advantage of the FosTRAP approach is that it circumvents the need for apriori knowledge of genetic markers and innervation patterns. This strategy could therefore be employed in future to gain a comprehensive understanding of individual nutrients, hormones, microbial metabolites, and inflammatory signals in gut-brain control of physiological processes and motivated behavior across health and disease states.

### Integration of post-ingestive reward by nigrostriatal circuits

Motivated behavior is driven by the ability to derive value from the outcome of an action. The nigrostriatal circuit, whereby SNc neurons release dopamine onto DS neurons, is essential for modulating acquisition and updating motivated behaviors ^66, 67^, including the motivation to eat ^21^. We and others have demonstrated that the presence of nutrients in the gut evokes dopamine release in the dorsal striatum. Importantly the amount of dopamine efflux correlates with caloric load ^18, 68^. Similar findings are also reported in human neuroimaging studies with striatal activity increased in response to food images ^69^. Furthermore, the use of fMRI and PET methodology demonstrate that the delayed post-ingestive response to food intake modulated dopamine signaling within the nigrostriatal circuit ^19^. Notably, pharmacological interventions that reduce dopaminergic tone have provided evidence for lower effort spending and motivation in both rodents ^22, 70^ and in humans ^71, 72^. Yet, these previous studies do not address if the decision to engage in motivated eating behavior is solely based on calories, or if type of calorie consumed is also encoded. We establish that separate nutrient-derived gut-brain circuits input into the nigrostriatal system to establish distinct motivation for fat or sugar. Foods that combine fats and sugars simultaneously engage two independent gut-brain circuits that reinforce the choice to select and consume nutritive foods.

Foods rich in fat and sugar are rarely found in nature ^73, 74^. Thus, prior to the development of industrialized farming in our recent past, the ability to locate foods with different sources of energy would have provided a competitive advantage to evolve macronutrient-distinct circuits for dopamine mediated learning and motivation. The modern food environment has led to the development of processed foods that incorporate high levels of fats and sugars ^75^. It has been previously demonstrated that human subjects want foods that combine fat and carbohydrate more than foods that contain fat or carbohydrate alone, and this was correlated with supra-additive striatal responses ^74^. Our results expand this prior work by identifying a neural mechanism to explain why the combination of fat and sugar is more rewarding than either alone. We find that mice voluntarily increase oral or intragastric intake of calorie-matched solutions that combine both fats and sugars compared to either fats or sugars alone suggesting that appetition occurs by post-ingestive signals independently of oral stimuli. These data are consistent with preference for oral stimuli (eg. flavor, texture, taste) being a conditioned response that predicts post-ingestive reward. This mechanism likely reflects a conserved system that evolved in rodents and humans to convey separate interoceptive information about the source of energy to the brain. However, in our current food environment, access to readily available foods rich in both fats and sugars results in greater valuation and appetition of obesogenic diets. Importantly, prior work demonstrates that vagal signaling is blunted in obesity ^76^, suggesting that interoceptive reward circuits increase the selection of calorically-dense and nutritionally-diverse foods and subsequently the susceptibility to diet-induced obesity, yet separate mechanisms are likely involved in maintaining a higher body weight set point.

## Conclusion

Our data support the idea that increased caloric intake of western diet results from hijacking interoceptive circuits that separately reinforce fats and sugars. Because interoceptive signaling occur below the level of consciousness, the motivation to consume obesogenic diets may occur without cognitive perception ^77^. In other words, conscious dieting efforts may be overpowered by the subconscious internal drive to consume foods rich in both fats and sugars. Therefore, inhibiting interoceptive gut-reward circuits may provide a viable therapeutic target for treating obesity by promoting voluntary reduction in the consumption of obesogenic diets.

## Methods Mice

All animal procedures followed the ethical guidelines and all protocols were approved by the Institutional Animal Care and Use Committee (IACUC) at the University of Florida (Protocol # 201810305) and Monell Chemical Senses Center (Protocol #s 1187 and 1190). Adult mice (6-20 weeks of age, both males and females) were used and maintained on a reverse 12-h light/dark cycle while housed at 22°C with *ad libitum* access to standard rodent chow (3.1 kcal/g, Teklad 2018, Envigo, Sommerset, NJ) unless otherwise stated. We did not observe significant sex differences between male and female mice in our experiments. All mice were on a C57BL/6J background, and transgenic genotypes validated by PCR. Strain details and number of animals in each group are as follows: C57BL/6J wild type: n=48: 24 male, 24 female; bred in house by UF breeding core, FosTRAP n=48: 24 male, 24 female; Jax B6.129(Cg)-Fos^tm1.1(cre/ERT2)Luo^/J (JAX stock no.021882), Fos Cre Tomato: n=48: 24 male, 24 female; bred in-house from Jackson Laboratory B6.129(Cg)-Fos^tm1.1(cre/ERT2)Luo^/J (JAX stock no.021882) and Ai14 (B6.Cg-Gt(ROSA)26Sor^tm14(CAG-tdTomato)Hze^/J, JAX stock no.007914), and Fos Cre Tomato Snap25: n=12, 6 male, 6 female; bred in-house from Jackson Laboratory B6.129(Cg)-Fos^tm1.1(cre/ERT2)Luo^/J (JAX stock no.021882), B6.Cg-Gt(ROSA)26Sor^tm14(CAG-tdTomato)Hze^/J (JAX stock no.007914), and Snap25-2A-GCaMP6s-D (B6.Cg-Snap25^tm3.1Hze^/J, JAX stock no.025111). Prior to experiments, mice were habituated for 2–3 days to experimental conditions such as handling, injections, attachment of gastric catheter infusion pumps, behavior chambers, attachment to patch cords for fiber photometry and optogenetics.

## Surgeries

### Nodose ganglia (NG) injection

Ten minutes before surgery, mice received a subcutaneous injection of carprofen (5 mg/kg; Henry Schein). Mice were anesthetized with 1.5-2.5% isoflurane, a 2 cm midline incision was made in the skin on the ventral aspect of the neck, underlying muscles, salivary glands and lymph nodes were retracted, and the vagus nerve was separated from the carotid artery by blunt dissection using fine-tip forceps. The NG was located by tracing the vagus nerve toward the head, and then exposed by retracting surrounding muscles and blunt dissection of connective tissues. A glass micropipette (30 μm tip diameter, beveled 45 degree angle) filled with either of the viral constructs (*pAAV9-Ef1a-double floxed-hChR2(H134R)-EYFP-WPRE-HGHpA* (Addgene 20298), *pAAV5-flex-taCasp3-TEVp* (Addgene 45580), *pAAV.Php.S-FLEX-tdTomato* (Addgene 28306), Addgene, Watertown, MA), CCK-saporin (ATS Bio, Carlsbad, CA), or saporin (ATS Bio, Carlsbad, CA) attached to a micromanipulator was used to position and puncture the NG. A Picospritzer III injector (Parker Hannifin, Pine Brook, NJ) was used to control injection speed and volume directly into the NG (total volume 0.5 µl). Incision was closed using 5-0 absorbable sutures in a running stitch and at 24 hours post-op analgesic was administered. Mice were fed with moistened chow and given at least 2 week for recovery and viral expression prior to experimentation.

### Retro Cre HPV application

In mice anesthetized with 1.5 - 2% isoflurane, analgesics buprenorphine XR 1 mg/kg and carprofen 5 mg/kg were administered subcutaneously 20 min prior to the surgery. An abdominal laparotomy, typically 1.5 cm, was performed. The liver was retracted to expose the HPV, and sterile parafilm was placed over the region with a fenestration over the HPV, to prevent spread of the viral vector. The viral retrograde vector (*pAAVrg-Ef1a-mCherry-IRES-Cre*, Addgene 55632; 1 µL per mouse) was applied with a small, sterile paintbrush to the HPV, brushing vigorously to ensure the viral penetration. Incision site was closed with 5-0 suture for the peritoneum, and wound clips for the skin and at 24 h post-op analgesic (carprofen) was administered. Mice were fed with moistened chow and given at least 2 weeks for recovery and viral expression prior to experimentation.

### Intragastric (IG) catheter implantation

IG catheters consisted of 6 cm silicon tubing (.047" OD x .025" ID, SIL047, Braintree Scientific, Braintree, MA) with 6 beads of silicon glue (#31003, Marineland, Blacksburg, VA) applied at 1 mm, 3 mm, 13 mm, and 15 mm from the distal end, and at 10 mm and 12mm from the proximal end, and a Pinport (Instech Labs, Plymouth Meeting, PA) for chronic intermittent access. In anesthetized mice, a 1.5 cm laparotomy was made, and the stomach externalized using blunt forceps and sterile cotton swabs. A 4 mm purse-string suture was made with 5-0 absorbable suture at the junction of the greater curvature and fundus, avoiding major blood vessels. The center of the purse-string was punctured with an 18-gauge needle, and the catheter is pushed in until the first drop of silicone glue is inside the stomach, the second drop exterior. The purse-string suture was then tightened and tied, anchoring stomach wall between the two drops of silicon glue. Next, a hole was punctured in the left lateral abdominal wall and fine-tip forceps used to pull the catheter through until the peritoneal wall wa between the third and fourth drops of silicone glue. A purse-string suture was made in the peritoneum around the catheter, tightened and tied to anchor the catheter to the peritoneum between the two drops of glue. The peritoneum laparotomy was closed with 5-0 absorbable suture. A 2 mm midline dorsal cutaneous incision was made caudal to the interscapular area, and a sterile blunt probe, 1 mm by 14 cm inserted and tunneled under the skin caudal to the left foreleg, to the abdominal incision. The catheter was threaded onto the end of the blunt probe and pulled through until the first bead on the proximal end is externalized, with the second proximal bead under the skin. The end of the tubing was cut off just above the external bead and a 22-gauge Pinport secured in the tube with superglue. A purse-string suture was made around the catheter, anchoring it to the skin between the two silicone beads. The abdominal incision was closed with suture clips. Analgesics buprenorphine XR (1 mg/kg) and carprofen (5 mg/kg) were administered subcutaneously 20 min prior to surgery and carprofen again at 24 h post-op. Mice were fed with moistened chow and given at least 1 week for recovery prior to experimentation. Daily body weight was monitored until pre-surgical weight was regained.

### In vivo 2-photon imaging

We used a 2-Photon microscope (Bruker) fitted with a Galvanometer for acquisition, and a piezo objective combined with a galvo/resonant scanner that allowed images to be obtained at a frame rate of 29 frames/second. The microscope was set up for *in vivo* imaging with a Somnosuite (isoflurane) anesthesia machine coupled to a homeothermic control warm pad (Kent Scientific) and a Programmable Syringe pump (Harvard apparatus PHD 2000, Cat no. 70-2002) for infusing nutrients in the gut.

Mice fasted for at least 2 h after the onset of dark were placed under continuous anesthesia (isofluorane/oxygen) and kept on a heating pad to maintain body temperature throughout the whole procedure. An incision (∼2 cm) was made above the sternum and below the jaw, the carotid and vagus nerve were exposed after separation of the salivary glands. Retractors were used to pull the sternomastoid, omohyoid and posterior belly of digastric muscle sideways and make the NG visible. The vagus nerve was cut immediately above the NG, which was carefully separated from the hypoglossal nerve and small adjacent branches. The vagus nerve was then dissected from the carotid and surrounding soft tissues, and the NG was gently placed on a stable imaging platform consisted of an 8 mm diameter coverslip attached to a metal arm affixed to a magnetic base. Surgical silicone adhesive (Kwik-Sil, WPI) was applied onto the vagus nerve to keep it immobilized on the coverslip and the NG immersed in a drop of DMEM (brand) media was covered with a second coverslip. Imaging was performed) using a 20x, water immersion, upright objective.

Infusions were performed with a precision pump that held syringes attached to a silicone tubing and filled with either sucrose, corn oil (Mazola) or saline. Prior to surgery to expose the NG, a small abdominal incision was made of the anesthetized mouse to expose the stomach. Next, the silicone tubing was inserted through a small incision in the stomach wall, into the proximal portion of the duodenal lumen. An exit port was created ∼2 cm distally to pylorus by transecting the duodenum. Super glue was applied onto the wall of the stomach to prevent the tubbing from sliding out of the intestine. To determine that there are distinct nutrient responsive subpopulations in the NG, neuronal activity of in response to different stimulus was monitored within the same mouse. Recordings of baseline neuronal activity signals started with continuous infusion of saline for 1 min (100 µl/min) followed by 5 minutes of 75% w/v sucrose infusion (100 µl/min), and then additional 10 min of saline flush (100 µl/min) to clear out the sucrose residues from the exit port. This was followed by the recording of a second baseline within the same animal after infusion of saline for 1 min (100 µl/min) followed by 5 min of corn oil (100%) infusion (100 µl/min).

GCaMP6s fluorescent changes were outlined in regions of interests (ROIs) with each ROI defining a single cell throughout the imaging session. The pixel intensity in ROIs (average across pixels) were calculated frame by frame (ImageJ) and exported to excel for manual analysis. The baseline signal was defined as the average GCaMP6s fluorescence over 5 min (fat or sucrose or saline) period prior to the stimulus introduction. Cells were considered responsive to nutrients if the following criteria were met: 1) the peak GCaMP6s fluorescence was two standard deviations above the baseline mean and 2) the mean GCaMP6s fluorescence was ≥ 70% above the baseline mean for a 20 second window around the peak. NG in which neurons did not present baseline activity were excluded from the study.

A Fos^TRAP^:Ai14 x SNAP25^GCaMP6s^ mouse line (n=4) was used to validate the selectivity of the FosTRAP approach (SupFig1A,B). Surgery and infusions were performed as described above. Briefly, saline infusion (1min, 100ul/min) was followed by infusions of equicaloric sucrose (75% w/v, 2 min, 100ul/min) and fat (33.3% Intralipid, 2 min, 100ul/min) solutions at different time points within the same mouse. An exit port of ∼5 cm was created distally to pylorus. For analysis, same criteria described previously was used and the response cut off was 20% above the baseline mean.

### Stereotaxic implants and viral injections

Mice were anesthetized mice with 1.5-2.5% isoflurane and placed in a stereotaxic apparatus (World Precision Instruments, Sarasota, FL) while resting on a heating pad. After skin incision (1-1.5 cm) and removal of all soft tissue from the surface of the skull, the periosteum was removed by blunt dissectionA dental drill was used to penetrate the skull above the target area. For viral GRAB^DA^ injections, a Hamilton Neuros syringe (Hamilton, Reno, NV) was filled with viral construct (1 µL, AAV9-hSyn-DA4.2, Vigene, cat. no. hD01) and lowered to the injection site in the dorsal striatum (anteroposterior (AP): +1.5, mediolateral (ML): ±1.5, dorsoventral (DV): -3.0) in a stereotaxic arm. A stereotax-mounted syringe pump (Harvard Apparatus, Holliston, MA) injected 0.2 µL per injection site at 100 nL/minute. Five min after completing the injection, the needle was slowly withdrawn, and skin closed with 5-0 absorbable suture.

For optogenetic and fiber photometry post implants, the protocol above was performed except additional holes were drilled (0.6-1 mm) nearby and stainless-steel screws secured allowing better fixation of the probe. Optogenetic posts were composed of fiber optic (FT200UMT, Thorlabs, Newton, NJ) secured in a ceramic ferrule (LC Zirconia Ferrule FZI-LC-230, Kientec, Stuart, FL) with UV-cured adhesive (RapidFix, St. Louis, MO). Posts were implanted over vagal terminals in the NTS (AP: -7.5mm, ML: ± 0.3mm, DV -5.0mm) and secured with dental cement (GC Fujicem 2, GC America, Tokyo, Japan). Fiber photometry optic fiber (400 µm core, NA 0.66, Doric, MF2.5, 400/430-0.) were implanted in the dorsal striatum (AP:+1.5, ML: ±1.5, DV:-3.0). Dental cement was used to cover the exposed skull, including screws, and allowed to set completely before mice were moved to the recovery area.

For microdialysis implants, the same protocol above was performed, except microdialysis guide cannulas (Amuza, San Diego, CA) were implanted in the DS as validated previously ^22^ (AP:+1.5, ML: ±1.5, DV:-3.0). An obturator or dummy cannula was used to keep the guide cannula patent, protruding at least 1 mm on the ventral side and the dorsal side screwed to the guide cannula thread.

## Behavioral Tests

### Food restriction

During all behavioral experiments, mice were maintained at 85-90% of body weight by restricting access to food. Briefly, for weight maintenance, approx. 2-3 g of food (10% of body weight) were given to mice 2 h after completion of the behavioral task. This paradigm ensured that mice learn to distinguish nutrients paired with the test solution without being confounding the association by any undigested chow in the gastrointestinal tract. In all cases, body weight was assessed every 24 h, just prior to refeeding. If any mouse weighed less than 85% of their starting body weight, they were fed 2.5 g plus the excess weight loss in g until they reached 85% of starting body weight again. *Ad libitum* water access was provided in home cage.

### Behavioral apparatus

All behavioral experiments - in conjunction with optogenetic stimulation and/or CNO infusions - were conducted in mouse behavioral chambers enclosed in a ventilated and sound attenuating cubicle (Med Associates Inc., St. Albans, VT). Each chamber was equipped with slots for sipper tubing equipped with contact lickometers with 10 ms resolution (Med Associates Inc.) used for licking detection. Separate chambers with a 5-hole nose poke wall with infrared beam detectors were used for nose-poke operant tasks (Med Associates Inc). These chambers were used in conjunction with optogenetic stimulation and/or pump-controlled infusion through a gastric catheter.

### Two-bottle choice flavor nutrient conditioning

Food restricted mice were placed in soundproof operant lickometer boxes (Med Associates) for training and habituation with one bottle of 0.2% w/v saccharin for 1 h per day for at least 3 consecutive days or until mice licked at least 1,000 times per h. Each day the bottle was placed in the right or left-hand sipper slot to avoid formation of a side preference. Once mice were trained to lick from the sippers, they underwent a pre-test, in which they were given 10-min access to two artificially sweetened (0.025% w/v saccharin) Kool-Aid (0.05% w/v) flavors in Med Associates lickometer boxes, one on the right and one on the left. At the 5-min point, the flavor positions were swapped to minimize the effects of individual side preference. The total number of licks for each flavor was tallied across the session and the nutrient (positive conditioned stimulus) was paired to the less preferred flavor and saline to the more preferred flavor to avoid a ceiling effect on preference for the nutrient-paired flavor. Flavor-nutrient pairings were counterbalanced within groups. After the pre-test, mice underwent 1-h conditioning sessions each day, receiving each intragastric solution with the paired flavor on alternating days for the following 6 days. To avoid the formation of a side preference during conditioning, mice received each flavor for an equal amount of time on each side of the box, alternating every 2 days. After 6 days of conditioning, on post-test day, the 10-min 2-bottle preference test was repeated, exactly as during the pre-test. Outcomes were measured by the percent preference each subject had for the nutrient-paired flavor during post-test.

### Nose-poke self-stimulation

Mice were trained to nose-poke for optogenetic stimulation in a Med Associates nose poke chamber with 2 nose-pokes, one active and one active following our previously published protocol. Mice were randomly assigned right or left active nose-pokes in a counterbalanced manner and underwent 3 days of 30-min nose-poke training during which a small amount of powdered rodent chow was placed in both nose pokes to motivate mice to explore the nose pokes. During all sessions, mice were tethered to a fiber-optic cable with swivel attached to laser and active nose-pokes resulted in optogenetic stimulation at 1 Hz frequency for 1 s duration at 5 mW intensity (measured at the fiber tip), as previously validated ^22^. Following 3 days of training, mice underwent 3 days of testing without food in the nose-pokes, during which the number of active and inactive nose pokes was measured over a 30-min test period each day.

### Microdialysis experiments

On the first day of microdialysis experiment, the cap of the guide cannula was removed, and a dialysis probe inserted into the guide cannula. After probe insertion, each mouse was placed in a microdialysis housing chamber and allowed to acclimate for 7 days with food and water available ad lib. Dialysate samples were then collected for up to 4 h while the animal was allowed to freely move around the cage. The dialysis probe was connected to an infusion pump by flexible Tygon tubing surrounded by a flexible spring tether attached to a liquid swivel and was perfused with artificial cerebrospinal fluid (aCSF) at a flow rate of 1 µl/min throughout the experiment. The tether was long enough for the animal to reach all parts of the test box unencumbered so that mice could comfortably eat, sleep or drink while attached to the tether. Sample collection continued during intragastric infusion of nutrients or after optogenetic stimulation and for about 3 h afterwards. Each animal received a maximum of 4 different stimuli. We performed 2 different tests per day with a 3-h interval between stimuli over a 3-day period, with one recovery between test days.

To assess DA release in the dorsal striatum during optogenetic stimulation of vagal fat or sugar responsive neurons, microdialysis measurements were performed in all subjects as described above. Specifically, during experimental sessions, microdialysate samples from freely moving mice were collected, separated, and quantified by HPLC coupled to electrochemical detection methods ("HPLC-ECD"). Briefly, a microdialysis probe (2 mm CMA-7, cut off 6kDa, CMA Microdialysis, Stockholm, Sweden) was inserted into the striatum through the guide cannula (the corresponding CMA-7 model). After insertion, probes were connected to a syringe pump and perfused at 1.2 µl/min with artificial CSF (Harvard Apparatus). After a 30-min washout period, dialysate samples were collected every 6 min and immediately manually injected into an HTEC-500 HPLC unit (Eicom, Japan). Analytes were then separated via an affinity column (PP-ODS, Eicom), and compounds subjected to redox reactions within an electro-chemical detection unit (amperometric DC mode, applied potential range from 0 to ∼2000 mV, 1 mV steps). Resulting chromatograms were analyzed using the software EPC-300 (Eicom, Japan), and actual sample concentrations will be computed based on peak areas obtained from 0.5 pg/µl DA standards (Sigma) and expressed as % changes with respect to the mean DA concentration associated with baseline samples.

## Histology

### TRAP protocol

In the TRAP protocol, mice were fasted 6 hours prior to intragastric infusion. 30 min before dark onset, mice received 500 µL intragastric infusate at 100 µL/minute in the home cage. As per previous literature ^78^, 3 h after the stimulus, mice were injected with 4-hydroxytamoxifen (4-OHT; 30 mg/kg, i.p.; MilliporeSigma, Burlington, MA), and normal chow returned to them 3 h after injection of 4-OHT. Intragastric infusates were 0.9% saline, sucrose solution (75%, 30%, or 15% w/v), an equicaloric fat solution (33.3%, 13.4%, 6.7% v/v; Microlipid, Nestle, Vevey, Switzerland), or an equicaloric combined solution of 50% fat and 50% sugar by caloric content.

### Perfusions

Anesthetized animals were transcardially perfused with phosphate buffer saline (PBS) until all blood was cleared, followed by 4% paraformaldehyde (PFA) in PBS for 2 minutes using the Leica Perfusion One System (Leica Biosystems, Wetzlar, Germany). Tissues including brain, NG, stomach, intestine, and HPV, were then harvested, post-fixed for 24 h in 4% PFA in PBS, before being transferred to 30% sucrose with 0.1% sodium azide for a minimum of 72 h before processing.

### Tissue processing & storage

#### Slicing

Whole brains were embedded in OCT and sliced in 1 in 3 series at 35 µm thickness on a Leica frozen microtome (CM 3050 S, Leica Biosystems) kept at -20°C. Slices were stored in cryoprotectant at -80°C until further staining or imaging. Nodose ganglia were embedded in OCT (Sakura, Torrance, CA) and sliced at 15 µM and frost mounted on Fisherbrand Superfrost Plus slides (Fisher Scientific) in 1 in 3 series and stored at -20°C until further processing.

#### Immunohistochemistry – cFos

Tissue was removed from cryoprotectant and rinsed 3 times for 10 min each in PBS at room temperature in 6-well plates, before being incubated for 30 minutes in blocking solution consisting of permeabilizing agent (244.5 mL PBS, 5 mL serum, 0.5 mL Triton-X100, 0.25 g Bovine Serum Albumin) and 20% normal donkey serum to prevent non-specific antibody binding. Tissue was then incubated overnight in permeabilizing agent with 1:1000 cFos antibody (Cell Signaling Technologies, rabbit-anti-cFos #2250) at room temperature. The next day, tissue was rinsed in PBS 3 times for 20 min each to remove unbound antibody, then incubated 2 h in permeabilizing agent containing 1:1000 secondary antibody (Donkey-anti-rabbit Alexafluor 647; AbCam, Cambridge, UK). Tissue was then rinsed 3 times for 2 h each in PBS, mounted onto plus slides, coverslipped with Prolong Diamond Antifade Mountant (Invitrogen, Waltham, MA), and stored at -20 °C until imaging and analysis.

### Tissue clearing

Tissue was cleared by incubating in TDE (2,2′-thiodiethanol, MilliporeSigma) for a minimum of 2 h prior to imaging.

### Imaging

Tissue sections labelled for cFos expression or mRNA probes were imaged using a Keyence BZ-X800 in a single plane with autofocus capture at 10x, or a Leica TCS SP8 confocal microscope at 20x. Quantification of cFos expression and colocalization was conducted using merged fluorescence images. For imaging of sections processed for in situ hybridization, positive and negative control probes were used to determine exposure time and image processing parameters for optimal visualization of mRNA signals and control for possible photobleaching. Brightness and contrast adjustment and automated quantification of cFos expression and colocalization was done with Nikon NIS Elements software.

#### Body composition measurements

Mouse body composition was measured with an EchoMRI-500 Body Composition Analyzer.

## Data Analysis

Data are presented as mean ± SEM and were analyzed for statistical significance as described in figure legends using Student’s t-test, one-way within-subjects or two-way within-subjects/mixed-model analysis of variance (ANOVA) (with the Greenhouse–Geisser correction applied as appropriate). Significant one-way ANOVA tests were followed by pairwise comparisons with Tukey’s correction for multiple comparisons. For two-way ANOVA, either simple main effects were reported, or significant interactions were reported and followed by pairwise comparisons with Sidak’s correction for multiple comparisons. The threshold for statistical significance was <0.05, and significant comparisons are reported in all figures as *p < 0.05, **p < 0.01, ***p < 0.001 and ****p < 0.0001. Comparisons in which p < 0.1 (but ≥0.05) are reported with exact p-values. Statistical analyses were performed in Microsoft Excel, or GraphPad Prism

## Acknowledgements

This study was supported by NIH grants R01 DK116004 (GL), R01 DK094871 (GL), and F31 DK1311773 (MM). We would also like to thank M. Afaghani at the University of Florida for technical assistance collecting pilot data.

## Supplemental figure

**Supplemental Figure 1.**
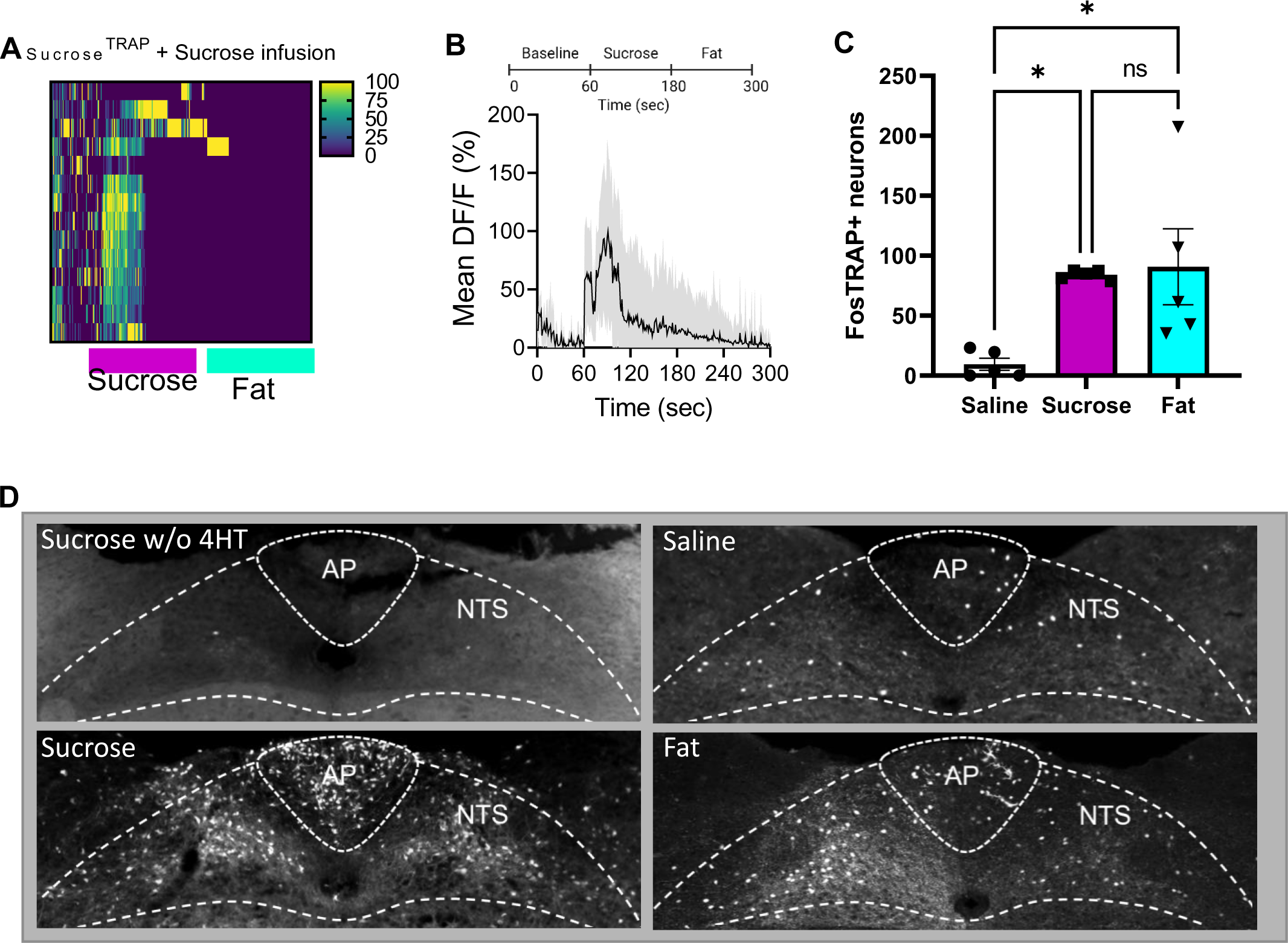
Validation of FosTRAP mice for genetic targeting of nutrient responsive neurons. A-B. In order to verify the specificity of the FosTRAP approach for targeting vagal sensory neurons we crossed FosTRAPxtdTomato mice to SNAP25-GCAMP6s. This allows us to visualize in red the neurons that are labeled with tdTomato during the TRAP process in response to intragastric infusion of sucrose, and record the real time activity patterns of all vagal sensory neurons using a mouse line that expresses the calcium indicator GCAMP6s expressed in all neurons. **A**, We find that the tdTomato neurons labeled in response to SugarTRAP have increased time-resolved responses (ΔF/F) in response to sucrose (15%, red bar, 2 min) with little activity in response to microlipid (7%, green bar, 2 min) infused into the proximal intestine. N=3 **B**, Average ΔF/F of GCaMP6s signals (normalized data) across all animals in Sugar^TRAP^ neurons increase in response to intestinal infusion of sucrose (60 to 180 s) but not fat (180 to 300 s). The dark lines represent means and lighter shaded areas represent SEM. **C-D.** To validate the FosTRAP approach centrally, tdTomato labeling in the NTS was assessed after intragastric infusion of saline (500µl/5min infusion), sucrose (15%), or microlipid (7%). **C**, Quantification of tdTomato in the NTS was significantly greater in response to sucrose or fat infusion.n=5. One Way ANOVA with Tukey post hoc analysis. **D** Representative images of tdTomato labeling in the NTS demonstrating elevated activity in response to sucrose or Fat compared to Saline, but absent in mice receiving intragastric sucrose in the absence of 4hydroxytamoxifen (4OHT) required for recombination of the inducible Cre. N=5. Data are presented as mean ± s.e.m. \**P* < 0.05, \*\**P* < 0.01, \*\*\**P* < 0.001, \*\*\*\**P* < 0.0001, NS, not significant.

**Supplemental Figure 2.**
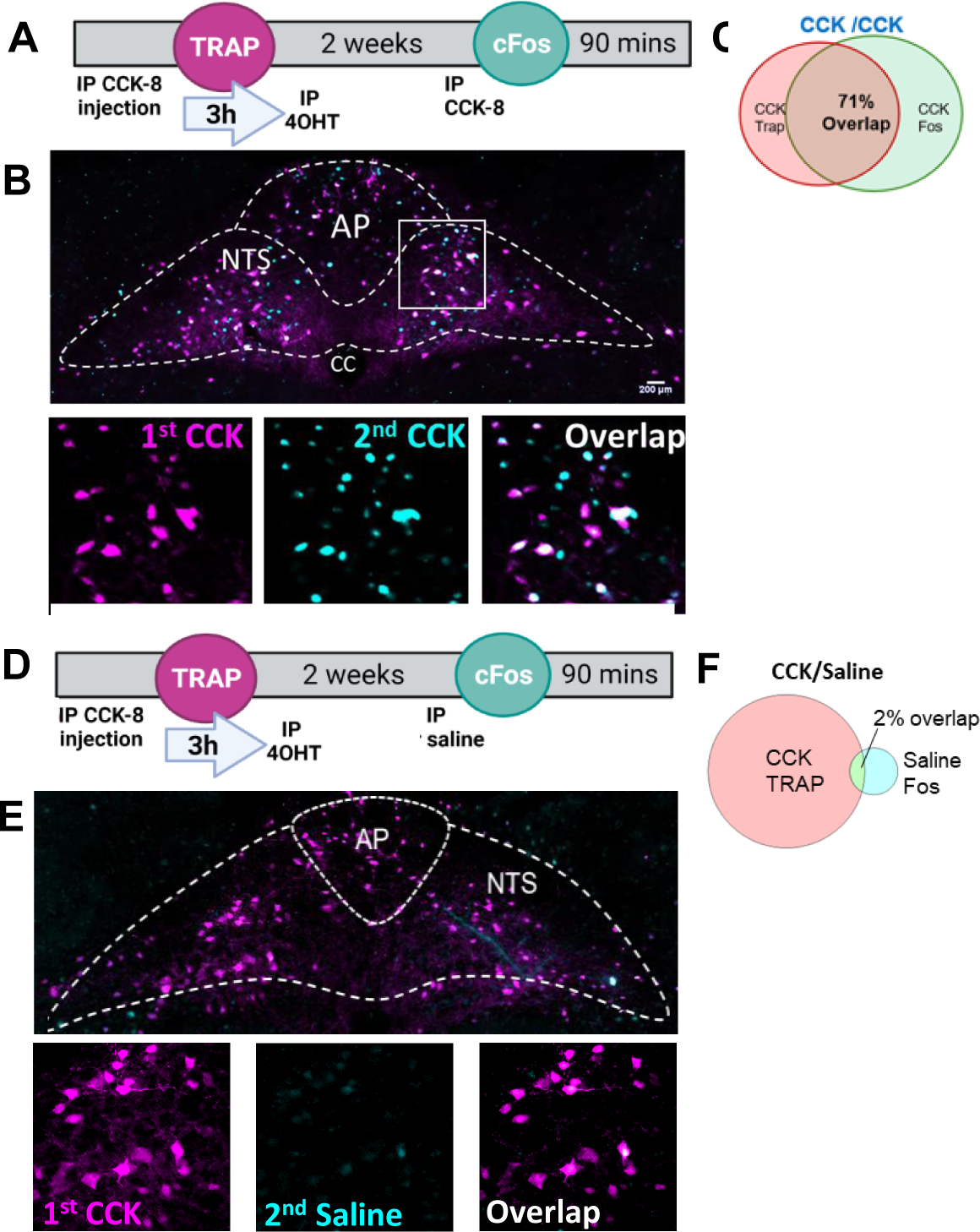
Validation of the FosTRAP model for comparing neural response to two different stimuli in the same animal. **A** To assess the reproducibility of the FosTRAP approach for comparing two peripheral metabolic stimuli at different timepoint within the same animal, we compared the overlap between tdTomato and cFos in response to two IP injections of the potent gastrointestinal hormone cholecystokinin (CCK8S, 4 µg/kg). **B** Representative images of the NTS in showing high overlap of active populations in response to two separate CCK injections administered two weeks. **C** Venn diagram showing high level of overlap. N=4 **D** Comparison of the overlap between an IP injection of CCK (tdTomato) followed by an IP injection of saline (cFos) two weeks later. **E** Representative images of the NTS demonstrating very low overlap between CCK and saline injections. **F** Venn diagram showing low level of overlap. N=4

**Supplementary Figure 3.**
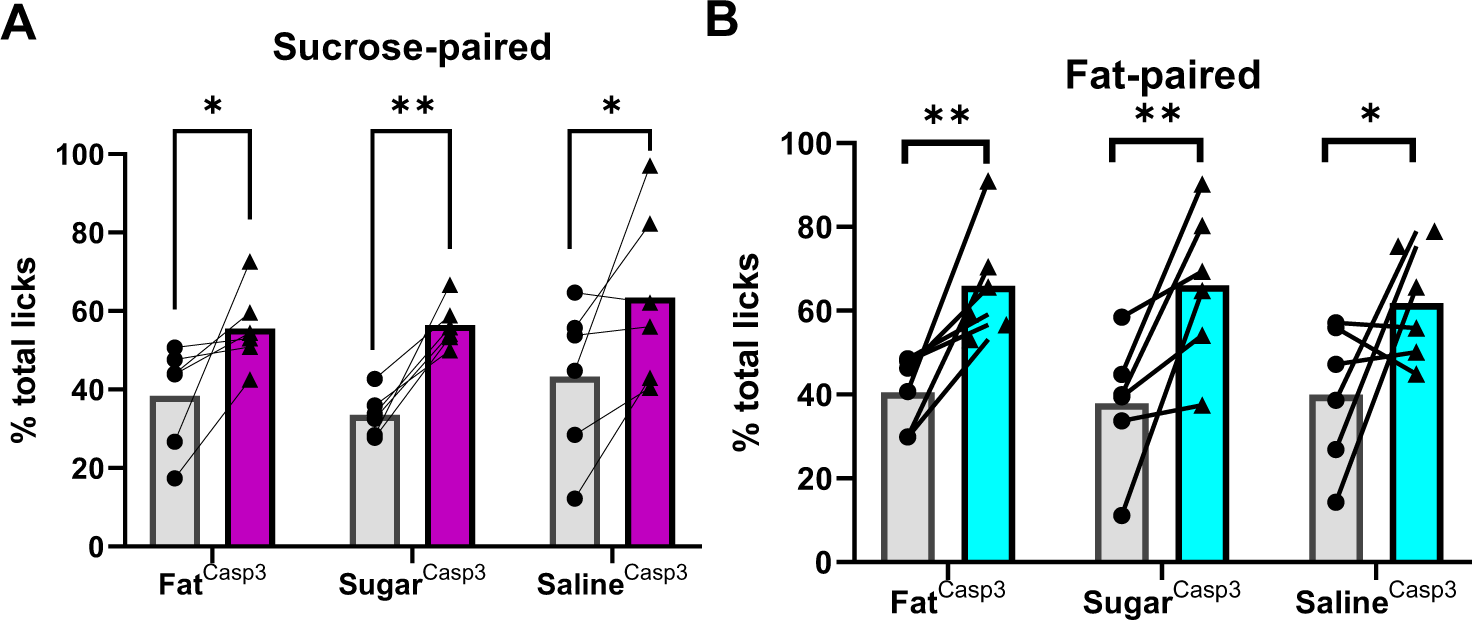
NG injections alone do not impair the ability of animals to learn nutrient-flavor conditioning. A-B. Control experiment for Figure 2F and G. Flavor nutrient conditioning was performed in all mice after bilateral caspase virus injection into the nodose ganglia but before trapping procedure to retain intact vagal signaling. All mice formed robust preferences to novel, non-nutritive flavors paired with either intragastric infusion of (**A**) sucrose or (**B**) fat before the targeted vagal ablation. N=6. Two Way ANOVA with Bonferroni post hoc analysis. Data are presented as mean ± s.e.m. \**P* < 0.05, \*\**P* < 0.01, \*\*\**P* < 0.001, \*\*\*\**P* < 0.0001, NS, not significant.

**Supplemental figure 4.**
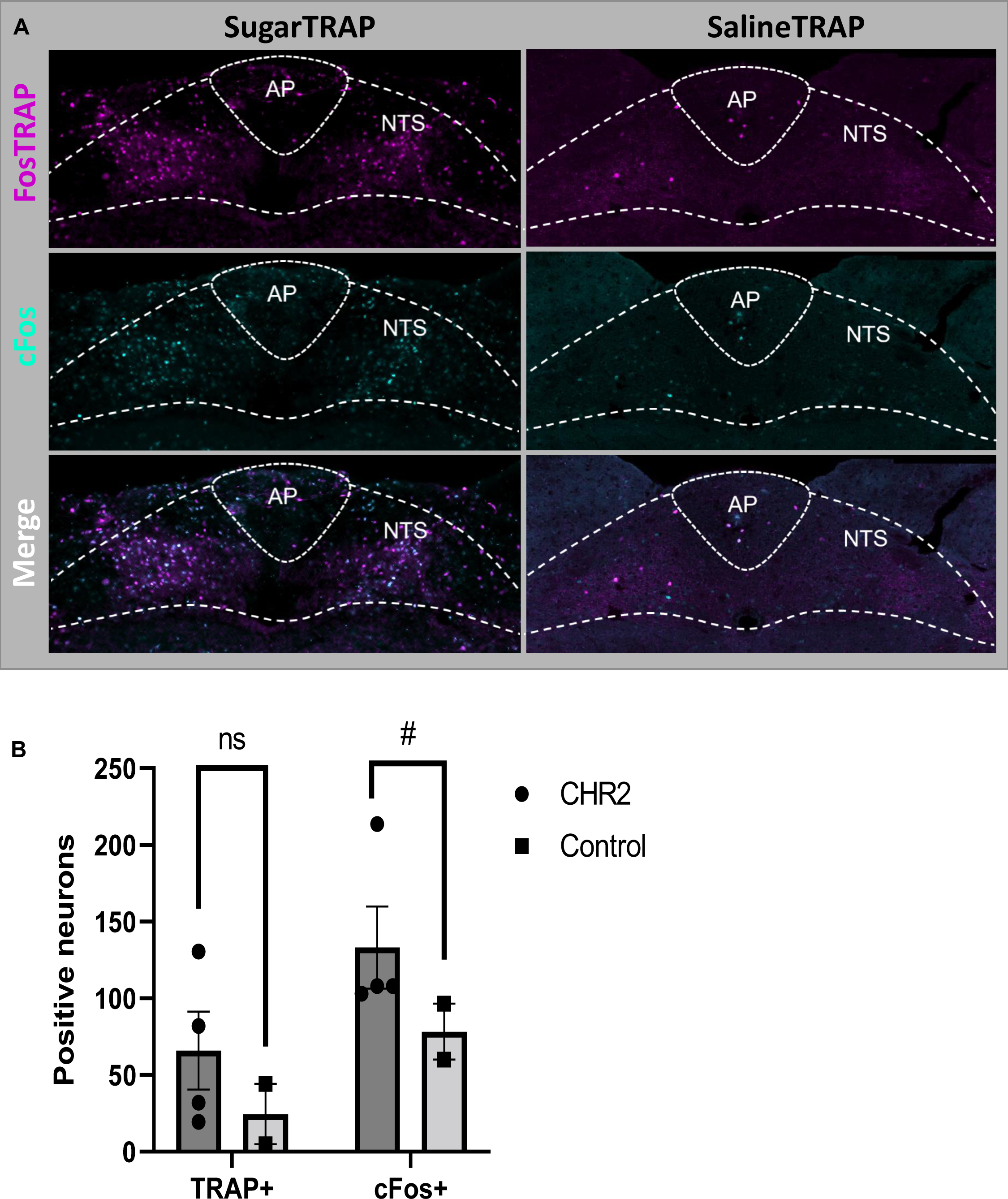
Validation of the optogenetic stimulation of NG. FosTRAP mice received bilateral injection of AAV9-EF1a-DIO-hChR2(H134R)-EYFP (ChR2) virus in the nodose ganglia and trapped in response to intragastric saline or sucrose infusions**. A** Representative images of cFos after laser stimulation was increased in the NTS of Sugar^TRAP^ mice compared to Saline^TRAP^ mice. **B.** The number of cFos+ NTS neurons was increased in response to blue light (473nm, 1 min/NG) stimulation in Sugar^TRAP^ and Saline^TRAP^ mice. N=2-4. Unpaired 2-tailed t-test. Data are presented as mean ± s.e.m. # *P* < 0.05, NS, not significant.

**Supplemental figure 5.**
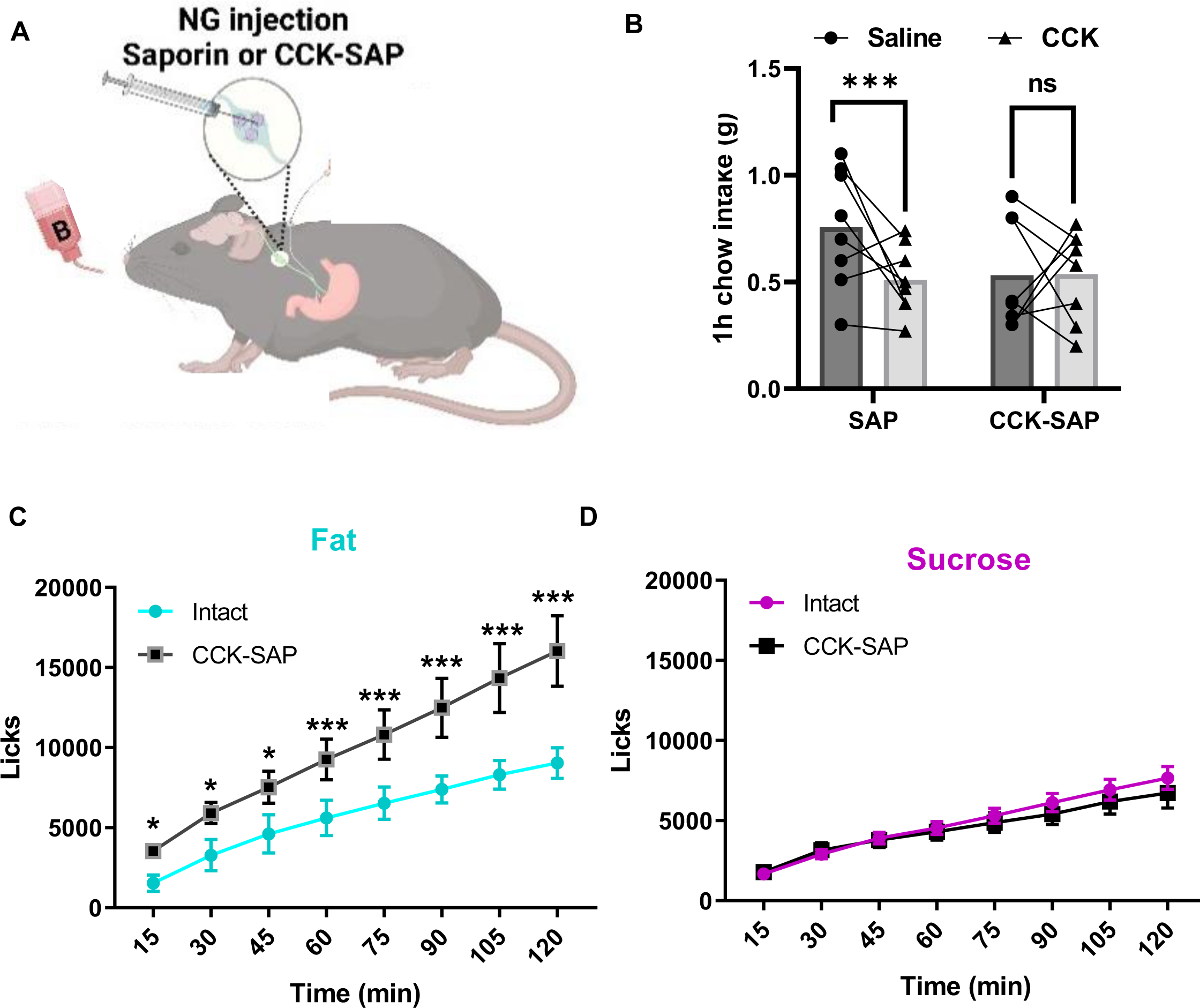
Vagal deafferentation of the gastrointestinal tract prevents fat but not sugar satiety. **A** Mice received bilateral injection of CCK-SAP or SAP into the NG. **B.** 1h chow intake was reduced in response to CCK (IP, 4 µg/kg) in control mice that were treated with bilateral NG injections of SAP, while CCK-induced satiety was abolished in CCK-SAP treated mice. N=8. Two Way ANOVA with Bonferroni post hoc analysis**. C.** Both SAP and CCK-SAP treated mice licked the same amount for sucrose solution (15%) presented in a bottle test. N=4. Two Way ANOVA with Bonferroni post hoc analysis**. D.** CCK-SAP treated mice significantly increased licking for microlipid solution (7%) compared to SAP treated mice. N=4. Two Way ANOVA with Bonferroni post hoc analysis. Data are presented as mean ± s.e.m. \**P* < 0.05, \*\**P* < 0.01, \*\*\**P* < 0.001, \*\*\*\**P* < 0.0001, NS, not significant.

**Supplemental figure 6.**
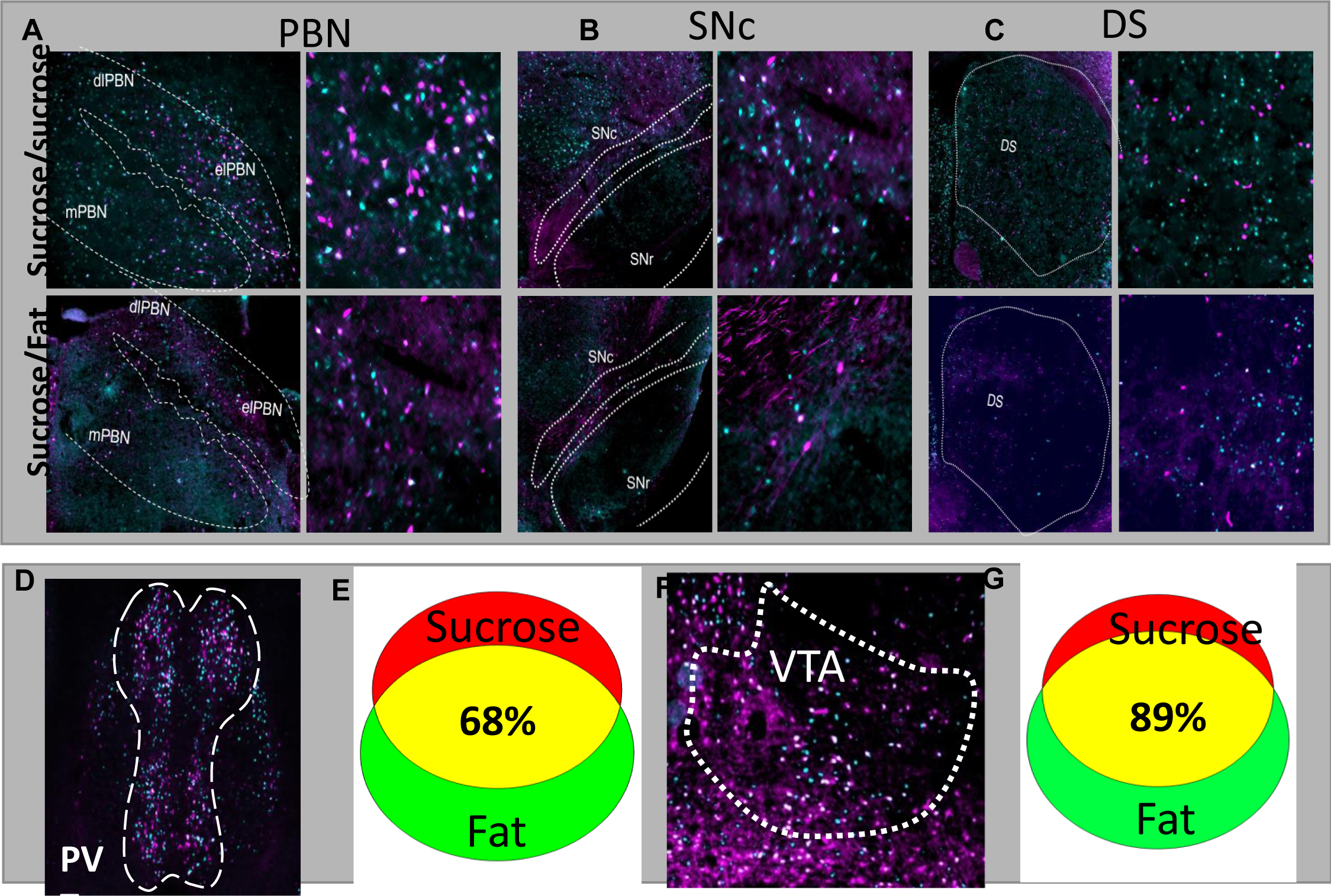
Fat and sugar recruit separate neuronal populations in the PBN, SNc, and DS, but highly overlap in the PVT or VTA. Data associated with Figure 4E comparing overlap between repeated IG infusions of sucrose or IG sucrose and IG fat in each brain region along the gut-reward circuit within the same animal. Low magnification on the left and higher magnification images on the right. A Representative images of the PBN show high overlap of two sugar infusions, but low overlap of sugar and fat infusions. B Representative images of the SNc show high overlap of two sugar infusions, but low overlap of sugar and fat infusions. C Representative images of the DS show low overlap in response to two separate infusions. D-G Not all brain regions discriminate between fat and sugar. D Representative images of the PVT showing high colocalization between neurons activated in response to intragastric sugar and fat. E Venn diagram quantifying colocalization from images in D. F Representative images of the VTA showing high colocalization between neurons activated in response to intragastric sugar and fat. G Venn diagram quantifying colocalization from images in F. N=3-7

**Supplemental figure 7.**
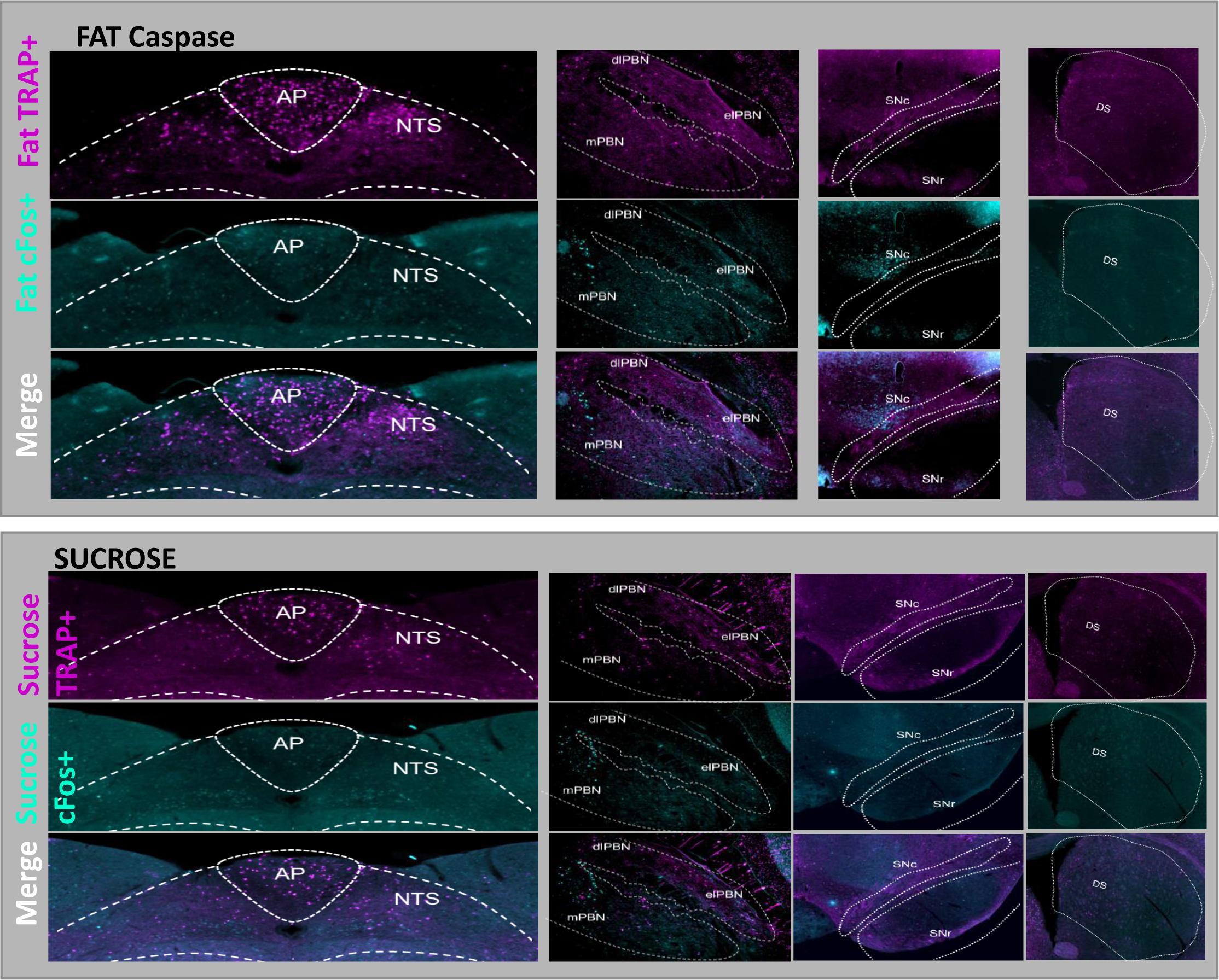
Nutrient vagal sensory neurons are necessary in recruiting the nigrostriatal reward circuits. Data associated with Figure 4H-J comparing overlap between repeated IG infusions of in the same mouse before (TRAP) and after (cFos) nutrient induced ablation of NG neurons. sucrose or IG sucrose and IG fat in each brain region along the gut-reward circuit within the same animal. **A** Representative images of fat responsive neurons in the NTS (1^st^ column), PBN (2^nd^ column), SNc (3^rd^ column), and DS (4^th^ column) before (magenta) and after (cyan) caspase ablation of fat sensing vagal sensory neurons. Low magnification on the left and higher magnification images on the right. **B** Representative images of sucrose responsive neurons in the NTS (1^st^ column), PBN (2^nd^ column), SNc (3^rd^ column), and DS (4^th^ column) before (magenta) and after (cyan) caspase ablation of sucrose sensing vagal sensory neurons. Low magnification on the left and higher magnification images on the right. N=3-6

